# Fragmented landscapes of mobile transposable element families bolster the diversity of large-effect alleles in Arabidopsis

**DOI:** 10.64898/2026.10.12.758581

**Authors:** Aurélien Petit, Erwann Caillieux, Mathilde Reignier, Ndeye Amy Gueye, Louna De Oliveira, Ciana Xu, Clio Cordier, Dersu Sezginmert, Vincent Colot, Pierre Baduel

**Affiliations:** Institut de Biologie de l’École Normale Supérieure, Université Paris Sciences et Lettres (PSL), UMR 8197 CNRS, U1024 INSERM, 46 rue d’Ulm, 75005 Paris, France

**Keywords:** transposable elements, pangenomics, Arabidopsis

## Abstract

Transposable elements (TEs) are powerful endogenous mutagens, but our understanding of the “mobilome” - the set of mobile TEs at the species level - has largely been shaped by the study of a single reference genome. To overcome this reference-centric view, we produced long-read genome assemblies for 91 *Arabidopsis thaliana* strains and performed a reference-free pangenomic graph analysis of structural variants. Our analysis revealed the presence of twice as many TE families with recent mobilization activity as previously described. These >200 newly identified TE families are ancestral, with a third experimentally shown to be mobile and although they are often characterized by the presence of full-length copies in only a few rather than in most strains, they nonetheless contribute significantly to the generation of large-effect alleles. These findings uncover a much more fragmented and impactful mobilome than previously documented and highlight how pangenome approaches dramatically improve the study of genetic diversity.

## Introduction

Transposable elements (TEs) are ubiquitous components of genomes and account for over 50% of total genomic DNA in many species, including humans and numerous crop plants ^1^. However, TEs capable of mobilization are only a very small fraction of the total TE complement of any given genome and our understanding of this mobile fraction, the “mobilome”, as well as of its mutagenic impact is far from complete. This is in part because many studies rely on a single reference genome and the annotated TE copies it contains. Although mobilome composition is expected to be highly similar across individuals in panmictic species, as is the case in *Drosophila melanogaster* ^2^ it is unclear how divergent it can be in species with strong population structure, such as in the model plant *Arabidopsis thaliana*.

A total of 15 mobile TE families have been experimentally identified in *A. thaliana*, most notably in the reference strain Columbia (Col-0) through the use of a mutation in the gene *DECREASE IN DNA METHYLATION 1* (*DDM1*), which encodes a chromatin remodeler that is key for the maintenance of DNA methylation and silencing of TEs ^3–11^. However, previous studies aimed at defining the *A. thaliana* mobilome at the species level using short-read sequencing data mapped against the reference Col-0 genome suggest that it is composed of about 150-200 distinct TE families, which belong to both class I (mainly Long Terminal Repeat or LTR-retroelements) and class II (DNA transposons), and the activity of which varies between strains ^12,13^.

With the advent of pangenome graphs based on long-read genome assemblies, it is now possible to identify structural polymorphisms in a reference-free manner, including the presence/absence TE insertion polymorphisms (TIPs) generated by transposition. Moreover, because these pangenome graphs contain the complete sequence of each TIP with precise boundaries, they can be used to create libraries of recently mobile TEs *de novo*. Here, we took advantage of the pantera tool, which was specifically designed for this purpose ^14^, to obtain a comprehensive and unbiased view of the *A. thaliana* mobilome based on genome assemblies we produced for 91 *A. thaliana* genetically diverse strains, including five for which we produced *DDM1* knock-outs. Our study reveals a much larger and highly-fragmented mobilome and illuminates how it fuels the generation of large-effect alleles at the species-level. More generally, our approach provides a novel and efficient reference-free framework for interrogating TE mobilization as well as its consequences in any species.

## Results

### Pan-genomic graph analysis reveals scores of additional, recently mobile TE families

In addition to 20 strains for which we previously produced a *de novo* genome assembly using Oxford Nanopore Technology (ONT) long-read sequencing ^15^, we selected another 71 strains that belong to the 10 main genetic groups identified in *A. thaliana* ^16^ (Fig. 1a) to best capture species-wide diversity and similarly produced a *de novo* genome assembly for each (Table S1). The maximum-likelihood phylogeny inferred from the coding sequences of shared BUSCO genes ^17,18^ using *A. lyrata MN47* (GenBank: GCA_944990045.1) as an outgroup confirmed genetic relationships established using SNPs (Fig. 1b, Fig. S1a). Furthermore, pairwise alignment (Fig. 1b) and alignment by chromosome ends (ABCE) (Fig. S1b) between the 91 genome assemblies indicated a high level of conserved synteny within chromosome arms and the presence of numerous large scale discontinuities in and around centromeres, consistent with previous findings ^19,20^.

**Figure 1.**
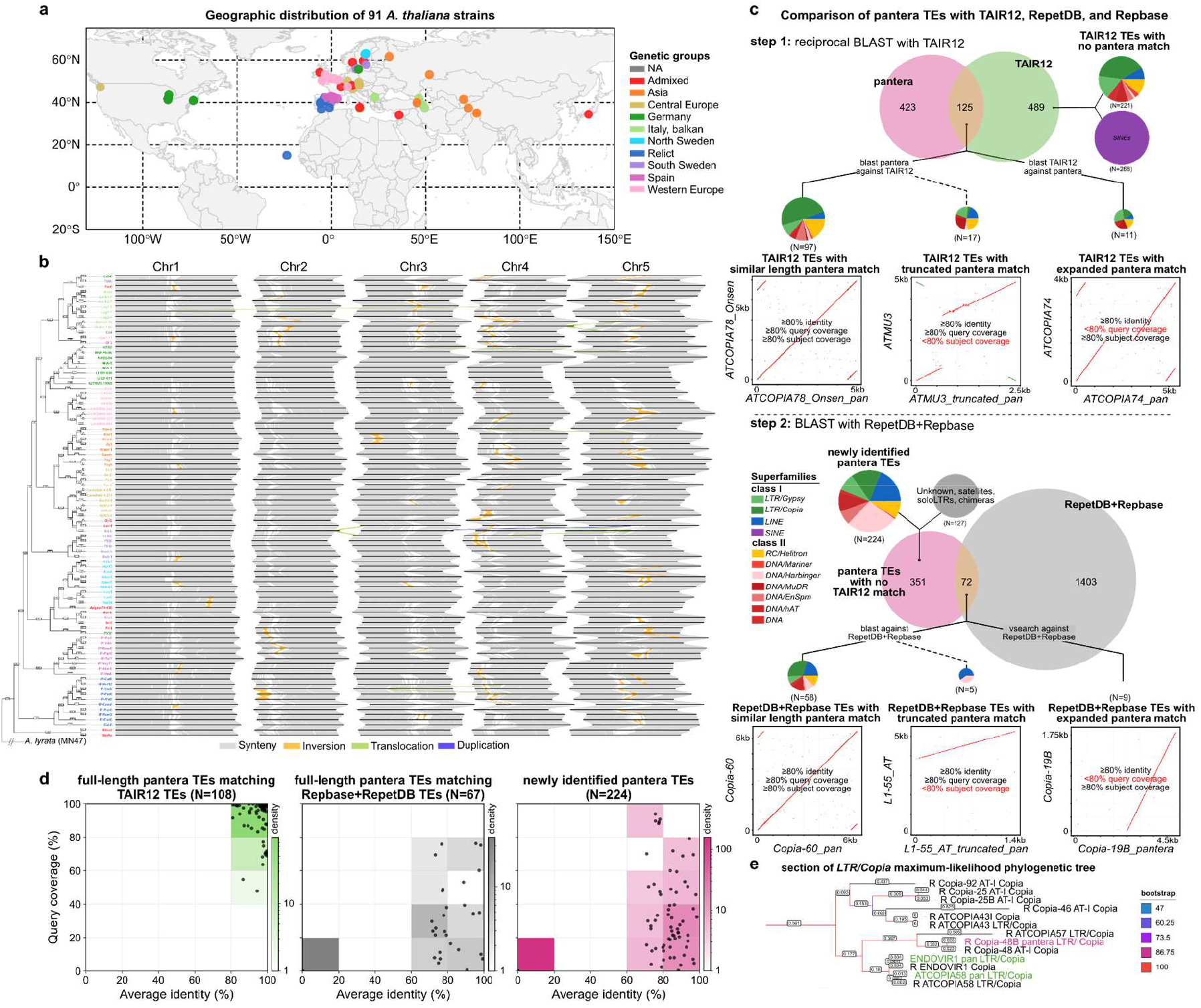
Detection of recently active TE families across 91 *A. thaliana* genome assemblies using the pantera tool. a) Geographic distribution of the 91 strains used, colored by genetic groups. b) Pairwise alignment of the 91 genome assemblies, ordered according to the maximum likelihood phylogeny built using BUSCO genes and with *A. lyrata MN47* taken as an outgroup. Conserved syntenic intervals are annotated in grey, and structural variants >500kbp are colored depending on their type. c) Comparison of the 421 pantera TE consensus sequences with the TAIR12 library of 602 TE sequences. As indicated below the Venn diagram, the 125 matches with 80% identity over 80% of sequence length (query or subject) fall into three categories, one with matches of similar length between TAIR12 and pantera and two with matches indicative of a single large truncation in pantera, or in TAIR12, respectively. For each category, its TE superfamily composition is indicated with a pie chart and a representative sequence dot-plot is shown as an example. The remaining 296 pantera sequences were then compared with the combined RepetDB TE library and Repbase TE database. The two categories of matches with the 80-80 rule we distinguished here are indicated, together with their superfamily composition and a representative sequence dot-plot, as above. d) BLAST results (with identity and query coverage on the x and y axes, respectively) against the TAIR12 TE library are shown for: pantera TEs that have a similar-length match in TAIR12 (green), or in RepetDB+Repbase (grey), or that have no match (pink). e) Part of the ML phylogeny for the *Copia* superfamily showing the relatedness of the TAIR12 and Repbase *Copia* sequences (black), with those identified by pantera. Non-truncated pantera sequences with a 80-80 rule match in either TAIR12 or RepetDB+Repbase are shown in green, and those with no such match are shown in pink. Branches are colored based on bootstrapping scores from IQ-TREE 2.

We then used pggb ^21^ (Fig. S1c) to build chromosome-by-chromosome pan-genome graph alignments between all 91 assemblies, and implemented pantera ^14^ to produce a *de novo* library of recently mobile TE families. After additional filtering to remove leftover redundancy (see Methods), we obtained 548 pantera TE consensus sequences (Table S2). Of these, 175 correspond, under the 80-80 rule (≥80% identity over ≥80% of the pantera TE), to TE sequences previously described, either in the TAIR12 TE library, or in the RepetDB TE library ^22^ and the Repbase database ^17^ (Fig. 1c-d). Apart from the 17 and five pantera sequences that exhibit a single large truncation compared to sequences from the TAIR12 library and RepetDB TE library or Repbase database, respectively (e.g. *ATMU3* and *L1-55_AT*; Fig. 1c), all others are full-length (e.g. *ATCOPIA78_Onsen* and *Copia-60;* Fig. 1c), including 12 of the 15 TE families with previous experimental evidence of mobility (Table S3) ^3–11^. In addition, through reverse BLAST against the TAIR12 TE library and vsearch clustering against RepetDB/Repbase (see Methods), we identified respectively 11 and nine pantera TE consensuses that match a sequence with a single large truncation compared to the pantera sequence (e.g. *ATCOPIA74* and *Copia-19B*; Fig. 1c-d).

Among the 351 pantera TE sequences with no matches, we removed 113 that were classified by RepeatClassifier ^23^ as Unknown, Satellites or as belonging to TE superfamilies not previously described in plants. We removed in addition four sequences that are likely solo-LTRs and upon manual curation another ten sequences that were found to be redundant or chimeric (see Methods). We thus obtained 224 high-confidence pantera TE sequences that represent newly identified TE families. In order to further explore the identity of these TE families, we built maximum-likelihood (ML) phylogenies at the TE superfamily-level using the TAIR12 library or Repbase database (see Methods and Supp. File 1). Using this approach, we could readily identify close relatives for 69 newly identified TE families, which we named accordingly (e.g. Repbase *Copia-48* and pantera *Copia-48B,* Fig. 1e, Table S2). As for the remaining 155 newly identified TE families, the majority belong to the *DNA/Harbinger, LINE, DNA/MuDR,* and *DNA/En-Spm* superfamilies, which were poorly represented in the TAIR12 TE library and for which we more than double their content (Table S2). Therefore, we named these 155 newly identified TE families simply in relation to the name of the corresponding superfamily (see Methods).

Together, these findings indicate that the diversity of mobile TE families at the species-level has been significantly underestimated so far. Indeed, most (489) of the 602 TE sequences present in the TAIR12 TE library that was derived from the latest version of the reference genome (https://github.com/oushujun/TAIR12-TE), including all 268 SINEs, were not identified by pantera (Fig. 1c) and therefore likely belong to fossil TE families, consistent with previous findings ^11^.

### Rare full-length copies of newly identified TE families reflect fragmentation of ancestral mobilome

To determine the genomic contribution of the TE families newly detected by pantera, we annotated each of the 91 genome assemblies using RepeatMasker (repeatmasker.org) and either the TAIR12 TE library or a ‘pan TE library’ (see Methods). This pan TE library (Supp. File S2) contains the 399 full-length consensus identified by pantera, including the 224 that do not match previously described TE families, as well as the 489 sequences of the TAIR12 TE library not found by pantera and the five RepetDB/Repbase sequences found as truncated by pantera (Table S4).

Annotation with the pan TE library covered on average ∼18Mb per genome, compared to ∼16Mb when using the TAIR12 TE library (Fig. 2a-S2a). Likewise, the number of full- or near full-length (>80% of consensus length, hereafter defined as full-length) TE copies detected with the pan TE library in each genome is also appreciably higher than with the TAIR12 TE library (+9% on average), notably in non-Germany strains (Fig. 2a-S2b). Furthermore, re-annotation of the TAIR12 genome assembly with the pan TE library also led to a noticeable increase (+13%) in TE content (Fig. 2b) and number of full-length copies (Fig. 2c). Moreover, although approximately half (283 out of 512) of the additional full-length copies were simply mis-annotated or truncated (see example in Fig. S2c-e) using the TAIR12 TE library, the other half (229) were fully or almost entirely missed (Fig. 2d; see example in Fig. S2f). Together, these findings demonstrate the higher sensitivity provided by the pan TE library, even for annotating the reference genome itself and its complement of potentially mobile TE copies.

**Figure 2.**
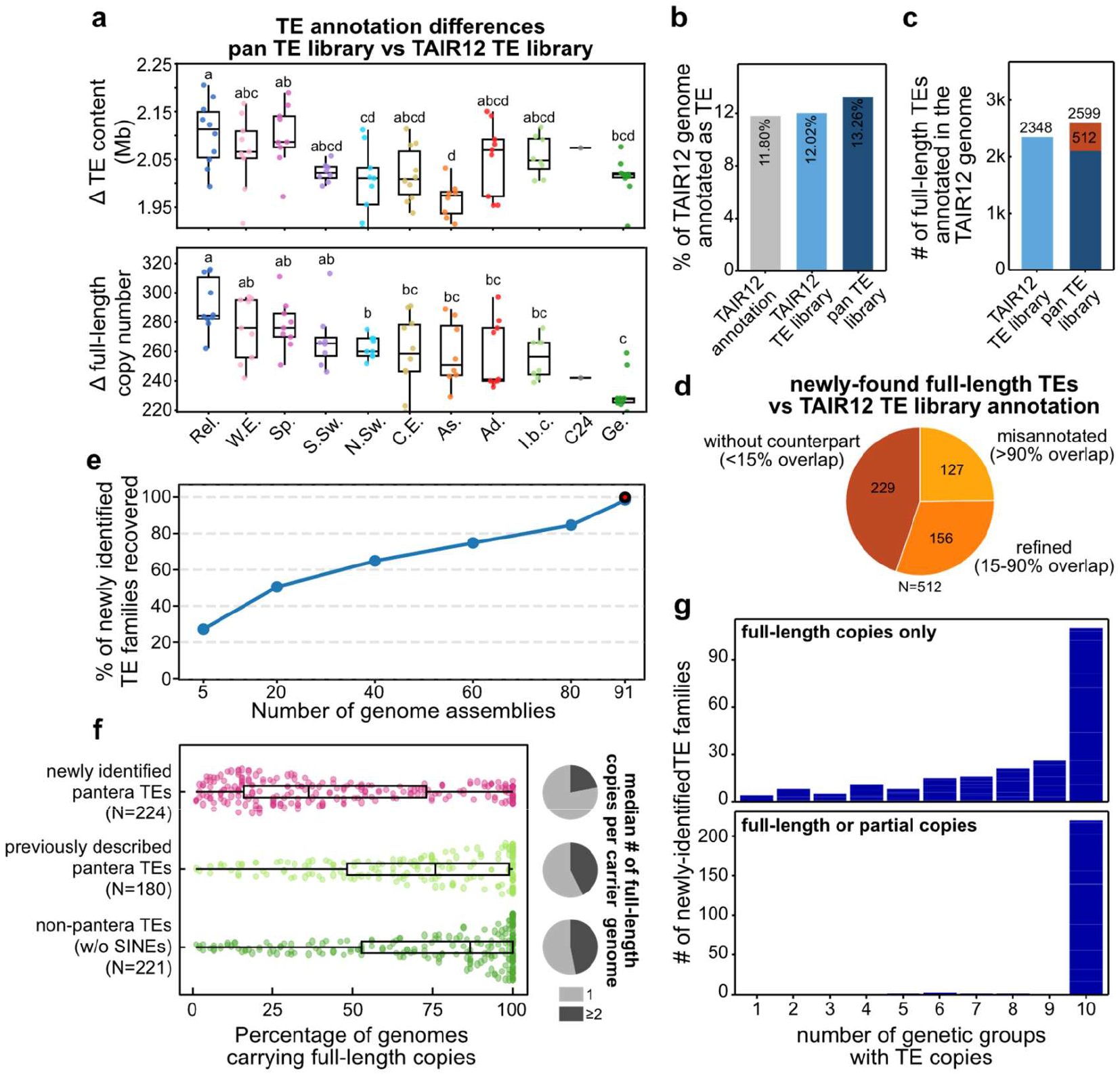
Fragmentation of the ancestral mobilome. a) Difference per genome in TE content (top) and number of full-length copies (bottom, excluding SINEs) based on TE annotation using RepeatMasker and either the TAIR12 or the pan TE library. Strains, with the exception of C24, are collapsed by genetic groups (Sp., Spain; Rel., Relicts; C.E., Central Europe; W.E., Western Europe; Ad., Admixed; Ge., Germany; N.Sw., North Sweden; As., Asia; I.b.c., Italy, balkans, caucasus; S.Sw., South Sweden) and letters above box plots indicate groups whose means are not statistically different from each other, based on post-hoc Tukey tests (see Methods). b) Comparison of TE content (in % of the TAIR12 genome) between the TAIR12 TE annotation, and the TE annotation of the TAIR12 genome assembly using RepeatMasker and either the TAIR12 or the pan TE library. c) Comparison of the number of full-length TE copies (excluding SINEs) based on TE annotation of the TAIR12 genome assembly using RepeatMasker and either the TAIR12 or the pan TE library. The 512 TE copies annotated as full-length exclusively with the pan TE library are indicated in red. d) Classification of these 512 TE copies in relation to their amount of sequence overlap with TE annotations obtained using the TAIR12 TE library. e) Percentage of the 224 TE families newly identified by pantera that are recovered using random subsets of 5, 20, 40, 60, 80 genome assemblies. f) Percentage of genomes containing full-length TE copies, for the three different categories of TE families: newly identified by pantera; previously described and also identified by pantera; not identified by pantera and represented in the TAIR12 TE library. Pie charts indicate for each category, the fraction of TE families that have on average (median), one or more than one full-length copy per carrier genome. g) Number of newly identified families per number of genetic groups with either full-length (top panel) or both full-length and partial (bottom panel) copies.

Despite these improvements, a fraction of the full-length TEs annotated by RepeatMasker using the pan TE library remain significantly divergent (>20%) from their corresponding consensus (Fig. S2g-h). This observation, combined with the fact that three of the 15 TE families with previous experimental evidence of mobility were not identified by pantera, presumably because of the small number of full-length copies they contain (Fig. S3a), indicates that our pan TE library does not capture the full extent of the mobilome diversity. Indeed, the number of TE families newly identified by pantera that are recovered when using fewer genome assemblies does not reach a plateau (Fig. 2e), which in turn implies that additional mobile TE families are likely to be discovered as more long-read genome assemblies and of better quality are produced^24^. This lack of saturation is also consistent with the fact that full-length copies of TEs belonging to the newly identified families (i.e. fulfilling the 80-80 rule with the consensus) are present only in a minority (30% on average) of genome assemblies (Fig. 2f). By contrast, full-length copies of previously identified TE families are typically present in 75-80% of genome assemblies on average, whether or not these TE families are also found by pantera (Fig. 2f-S2i). Furthermore, while genomes rarely possess more than one full-length copy in the case of the newly identified TE families, they tend to possess two or more such copies in the case of the TE families previously described (Fig. 2f).

Nonetheless, half (113) of the newly identified TE families have at least one full-length copy in each of the ten genetic groups, including relicts, and almost all (220) have at least either a full-length or partial copy (Fig. 2g). Moreover, among the 111 other newly identified TE families with a full-length copy missing from one or more genetic groups, 43 have a partial instead of a full-length TE copy at the same location in at least one of the missing groups (Fig. 2g, see Fig. S3b-d for an example). These findings indicate that most, if not all, newly identified TE families are ancestral and therefore do not represent cases of recent horizontal transfer. Indeed, despite the 5 million years of divergence ^25^ between *A. thaliana* and the two species *A. lyrata* and *A. halleri* (GenBank: GCA_964271285.1), RepeatMasker identified full-length, and thus potentially mobile copies in the reference genome of the latter two species for respectively 41 and 59 of the 224 TE families newly identified by pantera (Fig. S2j). This proportion is almost identical to that obtained for the 175 previously described TE families that are also identified by pantera, thus indicating a similar ancestrality for the two groups of potentially mobile TE families present in *A. thaliana*.

### Newly identified TE families generate large-effect insertion alleles

Previous work based on short-read sequencing data to detect TE insertion polymorphisms (TIPs) between strains revealed that the *A. thaliana* mobilome is particularly active ^12,13,26^, with an estimated transposition rate in nature that is at least of the same order of magnitude as the nucleotide substitution rate ^13^. Furthermore, many of the previously identified most mobile TE families exhibit strong insertion preferences near or within genes and thus tend to generate large-effect alleles ^11^, which are typically deleterious and therefore rapidly purged by natural selection ^13^. Given the many potentially mobile TE families newly identified or redefined by pantera, we set out to determine their contribution to the generation of TIPs. To this end, we relied on the GraffiTE pipeline ^27^, which uses a TE library to detect TIPs across the 91 genome assemblies and in relation to the reference genome. A total of 20,005 TIPs were detected using the TAIR12 TE library, almost all of which (98%) were also detected using the pan TE library, including 2,467 (13%) which were re-assigned to TE families newly identified by pantera (Fig. 3a; Supp. Data S1-2). Moreover, the latter library enabled the detection of an additional 4489 TIPs, the majority (66%) of which are contributed by TE families uniquely identified by pantera (2956 TIPs), although another 32% are contributed by previously described TE families also identified by pantera. We therefore conclude that the use of the pan TE library not only results in a higher sensitivity of TIP detection overall but also in a better assessment of the diversity of TE families contributing to the *A. thaliana* mobilome.

**Figure 3.**
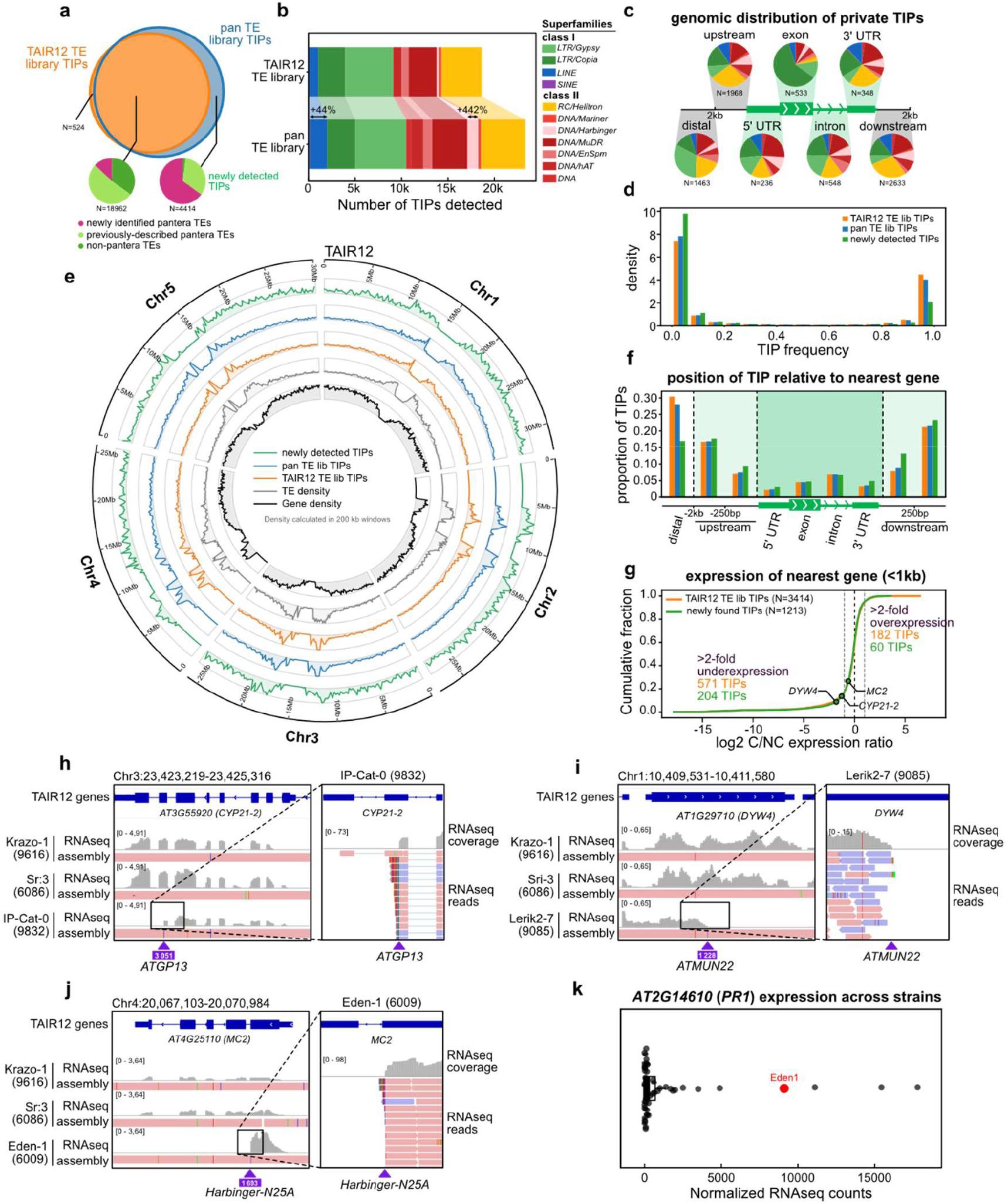
Extended diversity of large-effect TIPs generated by the newly defined mobilome. a) Overlap between TIPs detected by GraffiTE using the TAIR12 or pan TE library. Pie charts indicate the fraction of TIPs that are contributed by each of the three categories of TE families (see Fig. 2) and according to whether TIPs are detected exclusively when using the pan TE library or also when using the TAIR12 TE library. b) For each TE superfamily, number of TIPs detected by GraffiTE, using either the TAIR12 TE library, or the pan TE library. c) Private insertion alleles detected using the pan TE library, according to their genomic location and their TE superfamily identity. d) Frequency distribution of the TE-containing allele of the TE-containing allele of TIPs, depending on whether they were detected when using the TAIR12 TE library (orange), the pan TE library (blue) or exclusively using the latter (green). e) Circos representation of the density of genes and TEs along the five chromosomes of the TAIR12 genome assembly (TAIR 12 annotation) as well as of the three categories of TIPs shown in d. f) Proportion of TIPs detected using the TAIR12 TE library (orange), the pan TE library (blue) or exclusively using the latter (green), according to their location respective to genes. g) Cumulative distribution of expression fold change (log2) between carriers and non-carriers for genes with a nearby (<1kb) TIP, whether detected when using the TAIR12 TE library (orange) or exclusively when using the pan TE library (green). Numbers of TIPs associated with >2-fold over- or under-expression are indicated and fold-changes associated with the exonic insertion exclusively detected when using the pan TE library in the three genes *MC2, CYP21-2,* and *DYW4* are highlighted. RNA seq data were mapped on the TAIR12 genome assembly. h-j) Genome browser screenshots over the *MC2, CYP21-2,* and *DYW4* genomic regions of normalized RNA-seq coverage and genome alignments to TAIR12 for three strains, one of which contains the exonic insertion. Each insertion is associated with truncated transcripts, whose 3’ end extends to different degrees over the TE, as indicated by the short-read alignments shown on the right. k) Normalized expression of *PR1* across 52 strains with RNA-seq data, including Eden-1 (highlighted in red) which carries the *MC2* exonic insertion presumably responsible for high level of expression of *PR1* in this strain.

These improvements are particularly striking for TE families that belong to the *Harbinger* and *LINE* superfamilies, as we detected using the pan TE library respectively 442% and 44% more TIPs for these two superfamilies (Fig. 3b), which are both particularly poorly represented in the TAIR12 TE library (Fig. 1c). We therefore set out to examine the potential insertion preferences for these two superfamilies as well as for *Helitrons*, which were excluded from previous analyses because they do not produce TSDs upon insertion ^13^. To obtain a view that is least biased by genetic drift or natural selection, we considered, among all the TIPs detected using the pan TE library, only those where the TE insertion is present in a single strain (so-called private TIPs), as they are likely the most recent ^13^. Consistent with previous results ^13^, private TIPs produced by the *Copia*, *MuDR* and *Gypsy* superfamilies are enriched in exons, 5’UTRs, and away from genes (>2kb), respectively (Fig. 3c). In contrast, TEs belonging to two of the three previously under-studied superfamilies, namely *Harbinger* and *LINE,* show strong insertion preference near genes (≤2kb) and largely avoid exons and introns (Fig. 3c; S4a), highlighting their potential to generate regulatory variants and thus to enrich the palette of large-effect alleles produced by the mobilome.

Furthermore, the 4414 TIPs that are detected exclusively using the pan TE library are enriched in rare variants (Fig. 3d), and also in chromosome arms (Fig. 3e), especially near (≤2kb) or within genes (Fig. 3f), which suggest that previous analyses underestimated the mutational impact of the mobilome. Consistent with this interpretation, we detected using the pan TE library four TIPs, two previously described as well as two new ones, within the first intron of *FLC* (Fig. S4b), a key genetic determinant of flowering time in nature which is recurrently visited by TEs to generate large-effect alleles that can contribute to local adaptation ^12,13,28^.

To assess directly the functional impact of the TIPs that are detected exclusively using the pan TE library, we leveraged short-read RNA-seq data available for 52 out of the 91 strains ^29^, and compared the expression of genes within 1kb in strains with or without the insertion. For 1079 genes for which these comparisons could be made (see Methods), ∼24% display >2-fold changes of expression. Furthermore, in the vast majority of these cases (204 out of 264), expression is reduced in strains carrying the TE insertion (Fig. 3g). This result is consistent with previous observations ^13^ and also similar with those obtained when considering TIPs also detected with the TAIR12 TE library. Furthermore, visual inspection of some of the high fold-change TIPs within exons readily identified cases with transcript truncation at the position of the TE insertion, such as at the genes *AT3G55920 (CYP21-2)* and *AT1G29710 (DYW4)* (Fig. 3h,i). In addition, we noted an overaccumulation of transcripts corresponding to the first exon of *AT4G25110 (MC2*) associated with the presence of a *Harbinger* TE at the end of this exon in Eden-1 (Fig. 3j), mirroring that caused by a T-DNA insertion in the reference strain at approximately the same location (30bp upstream; ^30^. *MC2* encodes a metacaspase regulating pathogen-induced cell death^31^ and overexpression of its first exon alone was shown to induce a constitutive activation of immune responses, notably the central biotic response gene *PR1*, and to enhance pathogen resistance ^30^. Together with the finding that Eden-1 has among the highest levels of *PR1* expression of the 52 strains with RNAseq data (Fig. 3k), these observations indicate that the *Harbinger* insertion likely contributes to the very high levels of pathogen resistance observed in this strain ^32–34^.

### Fragmentation of mobilome activity is experimentally confirmed

Our pan-genomic analysis indicates that the *A. thaliana* mobilome is ancestral, but highly fragmented in its composition at the species level, because many TE families lack full-length, and thus potentially mobile copies in most strains (Fig. 2). To obtain direct evidence of this fragmentation, we used the dedicated tool TrEMOLO ^35^ (see Methods) to search for rare reads indicative of somatic or low frequency germline transposition in the ONT data we produced for the 91 genome assemblies. Using this approach and the pan TE library, we identified 509 so-called ‘outsiders’, i.e. TE insertions carried by one or two reads at most and with readily identifiable target site duplications (TSDs), except for *Helitron* TE families, which do not produce TSDs (Supp. Data S3). To obtain an independent assessment that outsiders correspond to bona fide insertion events, we tested 16 of them using PCR amplification and obtained confirmatory results for 13, the remaining three failing to amplify likely because of difficulties in primer design (see Methods and Table S5). Generally, outsiders are few (2-3 per 10X coverage) in a given strain, with noticeable exceptions (e.g. strain Qar8a, 26 outsiders per 10X coverage; Fig. 4a). The majority (54%) of outsiders belong to previously described TE families that are also identified by pantera, yet another 16% is contributed by newly identified TE families (Fig. 4b), consistent with these two distinct groups of TE families being active in many and few strains, respectively. Together, outsiders provide direct evidence of mobility for 141 pantera TE families including 59 of the 224 (26%) newly described TE families. In addition, we detected outsiders for 10% (51 of the 489) of the TE families not identified by pantera, indicating that, although mostly fossil, some of these TE families retain mobile copies, notably the *Helitron* family *ATREP1*, which contributed outsiders in 24 genomes in total.

**Figure 4.**
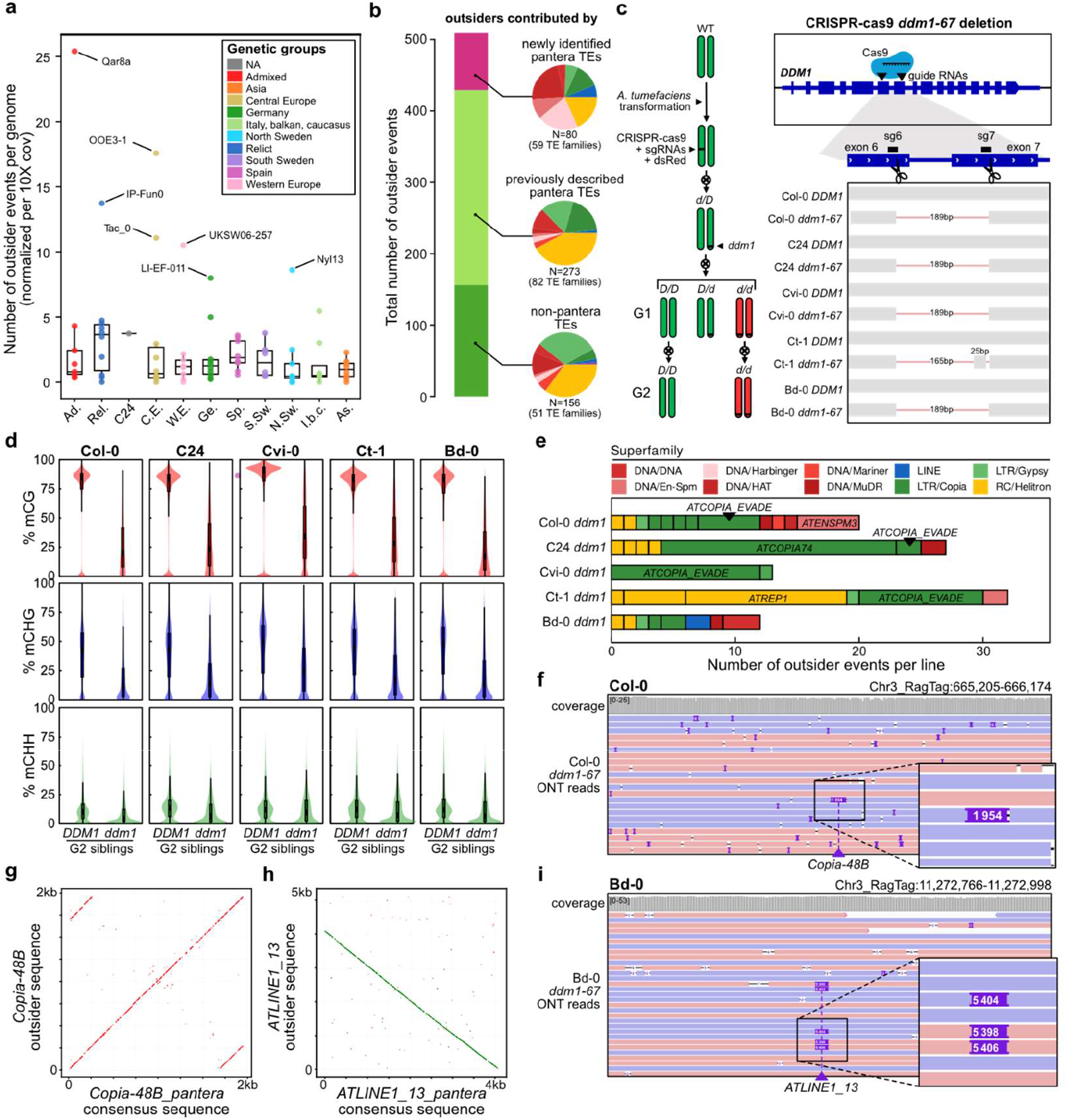
Experimental evidence of fragmented mobilome activity. a) Number of outsider events detected by TrEMOLO using ONT reads for each of the 91 genomes and normalized by 10X coverage. Results are shown by genetic group. b) Total number of outsider events contributed by TE families newly identified by pantera; previously described and also identified by pantera; and not identified by pantera. Pie charts indicate the relative contribution at the TE superfamily level. c) Schematics of the generation of the *ddm1-67* deletion allele using CRISPR-cas9 in 5 strains. Local sequence alignment indicating the size of the deletion in each strain. d) DNA methylation levels in the three contexts (CG, CHG, CHH) and over annotated TEs in homozygous second-generation *ddm1* mutant compared with their WT counterparts for each of the 5 strains. e) Number of outsider events detected by TrEMOLO using ONT reads of the five *ddm1-67* mutant lines. f) and g) Genome browser views of long-read alignments indicating the presence of an outsider in the mutant lines of Col-0 (*Copia-48B,* 1 read) and Bd-0 *(ATLINE1_13,* 5 reads). h) and i) Dotplots of the *Copia-48B* and *ATLINE1_13* outsiders against their pantera consensus.

In a second step, we used double-guided CRISPR-cas9 mutagenesis to generate a partial deletion allele of *DDM1* in four different natural strains (C24, Ct-1, Bd-0, Cvi-0) as well as in Col-0 for comparison (Methods and Fig. 4c), with the aim of compromising the epigenetic silencing of TEs in these strains and thus enhancing transposition activity. We verified by long-read sequencing that each homozygous mutant line (generation G2) we generated carried a large frame-shifting deletion starting in exon 6 and ending in exon 7 of *DDM1,* was devoid of the CRISPR-cas9 transgene and did not harbor any of the off-target mutations predicted using Cas-Offinder ^36^ (Fig. 4c-S5a). We named the knock-out allele *ddm1*-*67* in all five mutant strains, although the deletion harbored by the Ct-1 allele is slightly shorter than expected (165bp vs 189bp, Fig. 4c). Whole-genome bi-sulfite sequencing (BS-seq) indicated that in every case, plants homozygous for the *ddm1*-*67* mutant allele exhibit a significant loss of DNA methylation over TEs (Fig. 4c-d), consistent with previous results obtained in Col-0 with other *ddm1* mutant alleles ^7,37^.

Based on these results, we then used TREMOLO to identify transposition events in the five mutant strains (see Methods). Unlike in the WT, we considered here all outsiders carried by one or more reads so as to be able to capture germline events as well. Although this difference in filtering precludes a quantitative comparison of the two data sets, the large number of outsiders that we detected in the mutant lines (108 in total, contributed by 29 TE families; Fig. 4e; Supp. Data S4) is consistent with increased transposition activity as a result of the *ddm1-67* mutation. This observation notwithstanding, there were clear differences between mutant strains in the identity of mobile TE families. For instance, for *ATCOPIA_EVADE*, which is among the most transpositionally active TE family at the species level ^12,13^ as well as in lines derived from Col-0 *ddm1-2* ^11^ or *met1* ^38^, we detected numerous outsiders in the Col-0, Cvi-0, and Ct-1 *ddm1-67* lines (5-12 outsiders per line), but respectively only two and zero outsider in the C24 and Bd-0 *ddm1-67* lines (Fig. 4e). In contrast, there are numerous outsiders but exclusively in C24 for *ATCOPIA74* (Fig. 4e), whose longest copy is truncated in the Col-0 genome (Fig. 1c). Finally, we found outsiders for the newly identified *Copia-48B* family in the Col-0 background (Fig. 4f,h), where its full-length copy was misannotated previously (Fig. 2g), as well as for five other newly identified TE families, all in the Bd-0 background (see for example Fig. 4g,i). Altogether, these results demonstrate experimentally the mobility of one third of newly identified TE families as well as the fragmentation of the *A. thaliana* mobilome at the species-level.

### Evolutionary history of mobile TE copies explains mobilome fragmentation

To explore the genetic underpinnings of mobilome activity, we focused on the 108 transposition events we detected in the five *ddm1-67* mutant strains and searched for their donor copies. To this end, we extracted the sequence of each insertion and compared it, using either single-nucleotide pairwise diversity or phylogenetic relationships, to the full- or near full-length related copies that are annotated in the corresponding genome assembly. For 73 TE insertions results from both comparisons led to the identification of a single sequence, which designates it as the donor copy (see Methods). We identified in this manner 28 donor copies, which belong to 18 distinct TE families (Supp. Data S5). Remarkably, the donor copies with the highest number of outsiders tend to be the closest to genes (Fig. 5a), suggesting that, at least when the epigenetic silencing of TEs is compromised, gene proximity favors TE mobilization.

**Figure 5.**
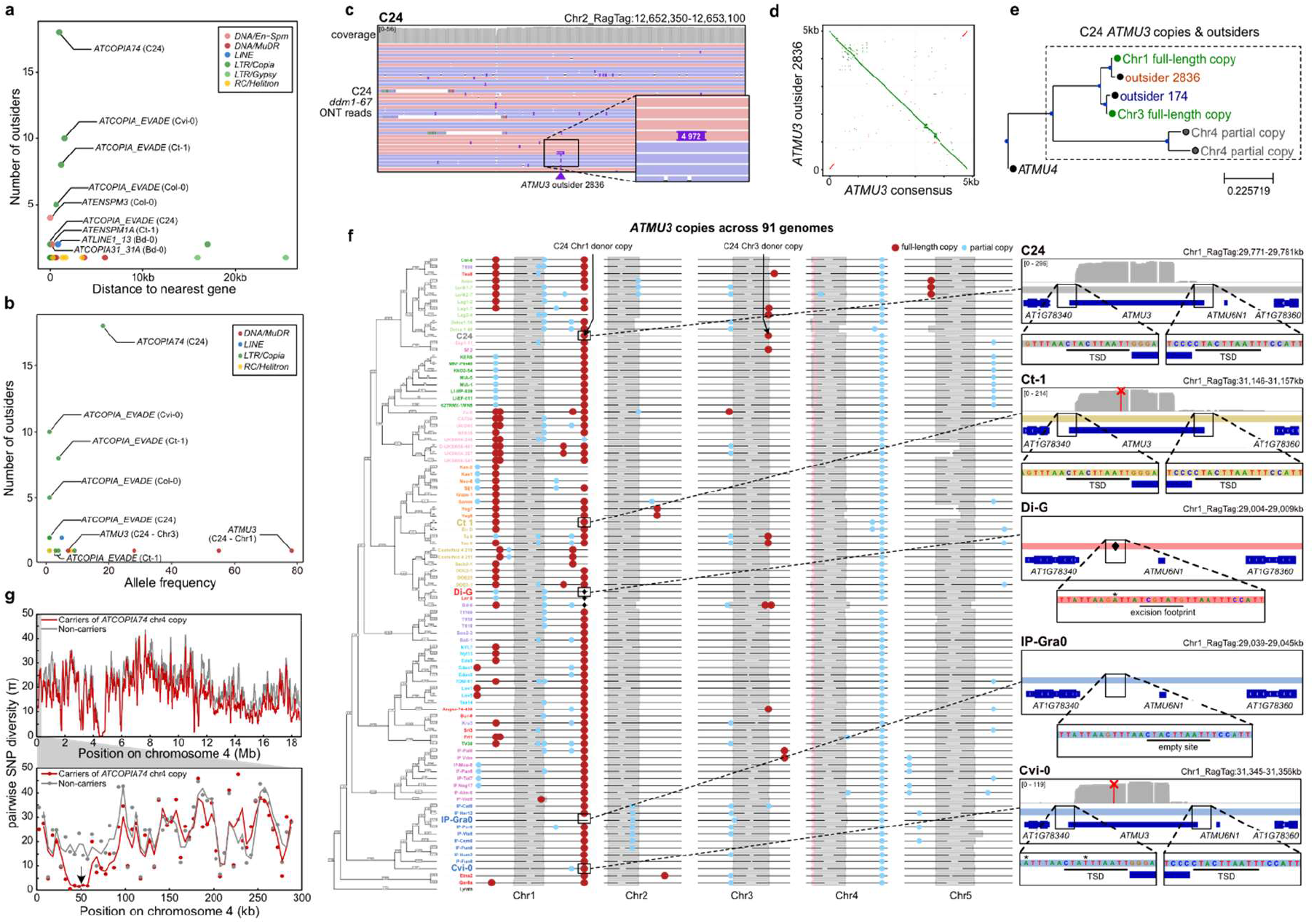
Contrasted evolutionary trajectories of mobile TE copies. a) Number of outsider events per donor copy in *ddm1-67* mutant lines in relation to its distance to the nearest gene.The identity of donor copies is indicated for those closest to genes. b) Number of outsider events per donor copy in the *ddm1-67* mutant lines in relation to the population frequency of that donor copy across the 91 genomes. The identity of donor copies is indicated only for those belonging to the *ATCOPIA_EVADE*, *ATMU3* and *ATCOPIA74* families. c) Genome browser view of long-read alignments indicating the presence of an *ATMU3* outsider in the C24 *ddm1-67* mutant line. d) Dotplot of the outsider sequence against the *ATMU3* TE family consensus. e) Neighbor-joining phylogenetic tree of *ATMU3* outsiders detected in the C24 *ddm1-67* mutant line (black dots) and of the *ATMU3* partial (grey) and full-length (green) copies the C24 genome, rooted using the *ATMU4* TE family consensus as an outgroup. Bar represents the proportion of nucleotide differences between sequences. f) Position of the *ATMU3* partial (blue) and full-length (red) copies present in the 91 genomes, aligned from their chromosome ends and ordered according to their phylogenetic relationship. Genetic groups are colored as in Fig. 1. The region carrying the donor copy of Chr1 identified in the C24 *ddm1-67* mutant line is highlighted on the right panels for five strains, which either carry the donor copy (C24, Ct-1 and Cvi-0) or an excision footprint (Di-G), or an empty site (IP-Gra0). Sequences at the insertion site are shown. TSDs, excision footprint, and empty site are underlined. Stars indicate SNPs in comparison with the empty site. Diamonds indicate excision sites. ONT long-read RNA-seq coverage is shown for the three strains for which we generated *ddm1-67* mutant lines (C24, Ct-1, and Cvi-0), together with the two distinct nonsense substitutions (red crosses) respectively present in the Ct-1 and Cvi-0 insertion. g) Pairwise nucleotidic diversity (π) across the TAIR10 chromosome 4 either whole (top panel) or within the first 300kb (bottom panel) for carrier or non-carrier strains of the full-length *ATCOPIA74* copy that is located at the beginning of chromosome 4 and which is mobile in C24. Lines represent moving averages of 20 (top panel) or 2 (bottom panel) consecutive 5kb windows, which are represented as individual dots. Arrow indicates the TAIR10 coordinates of the insertion.

For 19 donor copies, we could estimate their population frequency based on presence or absence of the insertion at the same locus in the other 90 genome assemblies (see Methods; Fig. 5b) and thus investigate their origin. Eleven donor copies were either private or present in just two to four genomes and the majority of these (6 donor copies) belong to the *ATCOPIA_EVADE* family (Fig. S6a), which is among the most transpositionally active in three of the five *ddm1-67* mutant strains (29 outsiders in total) as well as in a mutant strain of Tsu-0 deficient in CG methylation ^38^. Such a rapid turnover of mobile copies is that expected for TE families that tend to generate large-effect, mainly deleterious alleles ^13^.

In contrast to these 11 donor copies, the remaining eight donor copies have population frequencies that range from intermediate to high, with the highest frequency being that of the donor copy of the *ATMU3* family located on chromosome 1 and identified in the C24 mutant strain (Fig. 5b). Specifically, this donor copy, which produced one outsider (Fig. 5c,d,e), is found at the same location and also full-length or nearly so in another 77 strains (Fig. 5f). Furthermore, the 78 strains containing it encompass all 10 genetic groups and include three of the other four *ddm1-67* mutant strains (Col-0, Cvi-0, Ct-1) for which we did not detect *ATMU3* mobility. Also, visual inspection of the 13 strains that lack the *ATMU3* insertion identified the same excision footprint (Fig. 5f) in three closely related strains and a perfect empty site in the remaining 10 strains, which form an outgroup based on a phylogenetic analysis of the surrounding region (Fig. S6b, see Methods). Moreover, this local phylogeny mirrors that obtained genome-wide between the 91 genome and thus confirmed the ancient origin of the insertion rather than a widespread introgression across strains. Together, these findings demonstrate that the *ATMU3* insertion originated in the distant past, approximately 600 thousand years ago (kya; see Methods), with the empty allele representing the ancestral state and the excision allele a likely recent, secondarily derived state. Thus, TE insertions can retain mobility over hundreds of thousands of years, a conclusion that is at odds with the short residence time typically observed for TEs transpositionally active ^13^. Nonetheless, the *ATMU3* insertion has presumably lost its ability to transpose in the majority of genomes, as the coding-sequence of the long and abundant *ATMU3* transcript produced by the donor copy in the C24 mutant line (see Methods) is interrupted by premature stop codons in 55 strains, including the three strains for which we did not detect mobility in the *ddm1* mutant background (Fig. 5f).

Among the donor copies with intermediate population frequency, we investigated further that of the *ATCOPIA74* family, which is highly mobile (16 outsiders), but only in the C24 mutant strain (Fig. 5b). This donor copy is found at the same location on chromosome 4 as a solo-LTR in one genome (Lag2-4) and as a full- or near full-length insertion in another 16 genomes (Fig. S6c). Although these full-length copies are 99.6% identical, the one present in Bd-0 is truncated by a nonsense substitution in the sequence encoding its endonuclease domain, likely explaining why we did not detect mobility of this copy in the corresponding *ddm1-67* mutant line (Fig. S6c). Whatever the case may be, we estimated the insertion to have originated ∼40kya (max 6 SNPs over 5300bp), which is at odds with the fact that it is present in strains belonging to three genetic groups. To elucidate this observation, we carried out a phylogenetic analysis of the surrounding DNA sequence across all genetic groups as well as comparison of pairwise nucleotide diversity (Fig. S6d). Results revealed that the *ATCOPIA74* insertion belongs to a 22.5kb introgressed chromosome interval (Fig. 5g), which in turn indicates that it is part of a selective sweep, either directly or else as a result of genetic hitchhiking, another clear departure from the high-mobility / high-turnover model of TE mobilization.

Finally, although we could not estimate the population frequency of the donor copy of *Copia-48B* identified in Col-0, we noted that it is present at the same location in Cvi-0 (Fig. S6e), which suggests therefore that it is ancient, as it predates the event leading to the colonization of the Cape Verde islands by *A. thaliana* about ∼40kya ^39^. This last observation thus provides further evidence that TE insertions can remain mobile over significantly longer periods of time than previously thought ^13^.

## Discussion

In this study, we established a new pan-genomic framework, based on long-read genome assemblies and the dedicated pantera tool, to identify the nature and true extent of mobile TE families within species. Using this framework to investigate a panel of 91 *A. thaliana* genome assemblies that encapsulate the genetic diversity of the species, we uncovered the presence of at least twice as many recently mobile TE families as previously estimated ^12^. This result highlights the limited understanding of mobile TE families that one can achieve when analyses are mostly based on one reference genome only. Indeed, we discovered that many TE families possess full-length and thus potentially mobile copies in just a fraction of strains, thus preventing their proper identification using a single genome assembly. Moreover, we found no evidence that newly identified TE families reflect horizontal transfer, contrary to a previous suggestion ^20^. Rather, all newly identified TE families are at least as ancestral as the previously described ones, as they all similarly possess fossil copies in every genome analysed, including those of Relict strains. Moreover, many of these newly identified TE families have full-length copies in the genomes of the sister species *A. lyrata* and *A. halleri*.

Using the long-read sequencing data we produced for the 91 strains as well as for a subset of five strains for which we compromised the epigenetic silencing of TEs, we obtained experimental evidence of mobility for >150 of the newly and previously described TE families identified by pantera and confirmed the highly fragmented composition of the mobilome across strains.

Moreover, the evolutionary history of mobile copies revealed a broader diversity of TE dynamics beyond the typical high-activity high-turnover model of TE mobilization. At one end, we uncovered an ancient TE copy that maintained its ability to transpose for hundreds of thousands of years, while at the other end, we identified a more recent mobile TE copy that has been sweeping across distant genetic groups, as a result either of direct positive selection or else of genetic hitchhiking. Whether these are isolated cases or reflect a broader pattern of TE dynamics that are unaccounted for by current models remains to be established.

Thanks to the considerable expansion and better definition of the mobilome that resulted from the use of pantera, we uncovered thousands of previously undetectable TIPs, the majority of which are contributed by newly identified TE families. In fact, our findings indicate that these TE families, despite being each mobile in few strains only, are collectively a significant source of genetic variation at the species-level, further reinforcing the contribution of TE mobilization to mutation pressure in *A. thaliana* ^13^. Indeed, the TIPs produced by these newly identified TE families tend to have large effects, as they are often associated with expression changes of the genes near or within which they reside. Despite their functional importance, however, these TIPs, like other rare TIPs, are unlikely to be captured by forward genetic approaches such as GWAS, precisely because of their low frequency. This conclusion further highlights the necessity of properly capturing the diversity of the mobilome in order to fully characterize the genetic basis of natural phenotypic variation.

The extensive fragmentation of the *A. thaliana* mobilome stands in stark contrast with the uniformity reported for that of *Drosophila melanogaster* ^2^. This contrasted picture likely reflects the radically distinct demographics and mode of reproduction between the two species. While *D. melanogaster* is panmictic as a result of global movement through human trade, which explains the rapid invasion of this species by the *P* element and other TEs ^40,41^, *A. thaliana* is a sessile, autogamous species with limited outcrossing and strong population structure ^42^. The rapid accumulation of genome assemblies for numerous model and non-model organisms that are selfing or outcrossing, combined with the pan-genomic framework we report here, should soon provide a better understanding of the determinants of mobilome diversity and dynamics across species, and thus of the contribution of TEs to the generation of genetic diversity within each species.

## Methods

### Sequencing and *de novo* assembly of 91 *A. thaliana* genomes

Long-read ONT sequencing of 71 strains was performed as previously described for a first set of 20 strains ^15^. The contigs of all 91 genome assemblies were scaffolded using Ragtag ^43^ based on alignment on the recently published telomere-to-telomere genome assembly of the reference strain Col-0 ^44,45^.

To confirm the strain identity of each genome, Single Nucleotide Polymorphisms (SNPs) were called using NanoCaller ^46^ with “--mode snps--preset ont” options from the alignment of long reads on the TAIR10 reference genome generated by Minimap2 ^47^ with the “-ax map-ont” option. SNPmatch ^48^ was then used to obtain the best match across the strains of the 1001 Genomes.

For Principal Component Analysis (PCA), we used the short variants identified in the 1,001 Genomes (1001genomes_snp-short-indel_with_tair10_only_ACGTN.vcf from https://1001genomes.org/data/GMI-MPI/releases/v3.1/), and only kept bi-allelic SNPs with a minimum MAF of 0.05 and a minimum missing genotyping rate of 0.1. Plink ^49^ was then used to generate the PCA on the remaining SNPs.

We assessed assembly contiguity by calculating the contig N50 (Table S1) and assembly completeness using the embryophyta_odb10 database by Benchmarking Universal Single-Copy Orthologs (BUSCO v5.3.0) ^18^ with default parameters.

### Annotation of the 91 genomes

Gene annotations and pericentromeres were transferred from the TAIR10 reference genome ^50^ to each genome assembly using Liftoff ^51^.

### Phylogeny of the 91 genomes

Phylogenetic relationships were established by concatenating the alignments of the coding sequences of each of the 1,481 BUSCO genes shared across all 91 genomes and the *A. lyrata MN47* genome (GenBank: GCA_944990045.1). Alignments were produced using MAFFT ^52^ (--auto), trimmed using trimal ^53^ and concatenated using AMAS.py ^54^. A Maximum Likelihood tree was visualized with iTol ^55^, after building it using IQ-TREE 2 ^56^ with the model MFP+MERGE, based on the partition file and the supermatrix obtained with AMAS.py.

### Generation of pangenome graphs and use of pantera

We generated a pangenome graph for each of the five chromosomes using the 91 genome assemblies and the PanGenome Graph Builder (pggb) ^21^ version 0.6.0 with default parameters i.e. minimum nucleotide identity threshold of 95% (-p 95), a segment length of 10kb (-s 10000) and a k-mer size of 19 (-K 19). The five pangenome graphs were output in GFA format to be used as a single input for pantera.

We ran the pantera pipeline^14^ (https://github.com/piosierra/pantera), using the command “Rscript pantera.R -g gfa_list.txt -o output -d”, where the -d option avoids pairwise comparisons between genomes for faster processing and gfa_list.txt contains the paths to the pggb output in GFA format for the five chromosomes.

In order to remove residual redundancy from the pantera output, we performed clustering at 80% sequence identity using a combination of the greedy clustering algorithm vsearch --cluster_fast (--id 0.8 --strand both --iddef 2) and BLAST, using the raw pantera output as both query and subject. Non identical and non-overlapping matches that cover at least 80% of the length of any pantera sequences, with an average identity above 80%, were clustered together. From each cluster, the longest non-tandem TE sequence was then extracted. This step eliminates redundancy directly from the pantera output, notably solo-LTRs, reducing the number of sequences from 958 sequences to 548.

### Comparison of TE consensuses obtained using pantera to existing TE libraries and database

First, we compared the 548 TE consensuses obtained using pantera to the sequences of the TAIR12 TE library (https://github.com/oushujun/TAIR12-TE, version 30/07/25). Given that this library keeps separately the LTR and INT sequences of LTR retrotransposons, we started by stitching back together the LTR and INT sequences belonging to the same element. LTR and INT with no match were kept as standalone. After this stitching step, we performed the same clustering approach as for the pantera raw output (see above) and removed in this manner 20 redundant sequences from the TAIR 12 TE library. Matches between the 548 pantera TE consensuses and the 602 sequences we obtained after curation of the TAIR12 TE library were identified using BLAST and the pantera TE consensuses as queries. A pantera TE consensus was deemed to match a sequence from the TAIR12 TE library when, after concatenating all the non-overlapping hits returned by BLAST for that TAIR12 sequence, the concatenation covered, with an average percentage of identity >80%, at least 80% of the length pantera TE consensus. Using this approach, 97 pantera consensuses were found to have a full-length match in the TAIR12 TE library, and another 17 pantera consensuses were found to have a match, but only over less than 80% of the length of the matching sequence in the TAIR12 TE library. Dotplot analyses (https://en.vectorbuilder.com/tool/sequence-dot-plot.html) indicated that all of these 17 pantera consensuses present a single large truncation compared to their TAIR12 counterpart. Using a reciprocal BLAST (sequences of the TAIR12 TE library as queries), we identified a further 11 pantera consensus with a match in the TAIR12 TE library, but this time over less than 80% of the pantera sequence. Dotplot analyses indicated that in each case, the matching sequence of the TAIR12 TE library presents a single-large truncation compared to the pantera consensus.

Second, we compared the 423 pantera consensuses with no match in the TAIR12 TE library to sequences of the RepetDB TE library ^22^ and the Repbase database ^17^, using BLAST and the pantera consensuses as queries, and applying the 80-80 rule, as above. We identified in this manner full-length matches for 58 additional pantera consensuses. Another five pantera consensuses have instead a match, but only over less than <80% of the sequence present in the RepetDB TE library or Repbase database. Dot plot analyses indicated a single large truncation of the pantera sequence compared to that of its match in all five cases. Finally, by using BLAST and vsearch --cluster_fast (--id 0.8 --strand both --iddef 2), we identified nine other pantera consensuses that have a matching sequence in RepetDB or Repbase, but this time with the latter presenting a single large truncation compared to the pantera consensus.

Thus, we identified matches for 197 pantera consensuses in total, 22 of which present a truncation compared to their matching sequence, and no matches for 314.

### Curation of the pantera consensuses

The 314 pantera consensuses with no matches were classified using RepeatClassifier ^23^ and after manual curation to remove consensuses classified as Unknown, Satellites, LTR-retroelements (*Gypsy* or *Copia*) less than 900bp-long, which likely correspond to left-over solo-LTRs, or *LINE/Penelope, LINE/L1-Tx1, DNA/Novosib,* and *LTR/ERV1,* as these TE superfamilies were not previously identified in plants, we obtained a set of 234 pantera consensuses corresponding to newly identified families.

These 234 pantera consensuses as well as the 175 pantera consensuses with similar or longer length than their match in the pre-existing libraries or database were then curated using TEtrimmer ^57^ with the option: --min_seqs 2 on a sample of 10 genomes that represent all 10 genetic groups. For TEs that were too low-copy in this sample of 10 genomes, TEtrimmerGUI was used manually. This curation step helped improve 97 pantera consensuses, most of the time because the use of the dual (-d) mode in pantera can lead to poor resolution of complex structural variants, such as nested TEs. In addition, five consensuses were found to be identical to another five, which were thus removed and another five consensuses did appear highly chimeric and were also discarded.

### Creation of a pan TE library

We created a pan TE library comprising the curated 175 full-length pantera consensuses with matches, the curated 224 pantera consensuses with no matches, the five Repbase sequences with a truncated pantera match, and the 489 TE sequences of the TAIR12 TE library with no pantera match. Therefore, the pan TE library contains 893 sequences, almost half of which are composed of high-confidence and curated pantera consensus.

### Phylogenetic analysis of the sequences composing the pan TE library and naming scheme

The 893 sequences of the pan TE library were regrouped and aligned by superfamilies with MAFFT (--reorder --adjustdirectionaccurately) ^52^ in order to build Maximum Likelihood phylogenetic trees. Trees were visualized on iTol ^55^ based on IQ-TREE 2 ^56^ using the model -m TEST and -bb 1000. The 175 full-length pantera consensuses with matches were given the name of the matching sequence and adding the suffix “_pan” (e. g. matching TAIR12 name: *ATCOPIA78_Onsen*; new name: *ATCOPIA78_Onsen_pan*), to distinguish them from the 489 TE sequences of the TAIR12 TE library with no pantera match. As for the 224 pantera consensuses with no match, we named them based on the closest known sequence and adding “_pantera” (e. g. closest sequence: *ATCOPIA5*; new name: *ATCOPIA5B_pantera*), except when that sequence was too distant or when bootstrapping did not provide strong support for the branching, in which case the pantera consensus was named simply using the name of the superfamily and adding as a suffix the smallest unused number followed by “_pantera” to distinguish them from the others (e.g. the last Gypsy name given in the TAIR12 or RepetDB TE library or in Repbase database is *ATGP11*, therefore: *ATGP12_pantera* for Gypsy).

### Alignment by chromosome ends of the 91 genome assemblies

Genome alignments were carried out from both ends of chromosomes to accommodate the large variations in centromere length and structure that can exist between strains for the same chromosome ^58^. To this end, we first determined, for each of the five chromosomes, the longest assembly across all strains. To standardize chromosome lengths, we introduced in each of the other chromosome assemblies a gap compensating the difference in length at the position of the contig breakpoint (stretches of 100 Ns produced by RagTag ^43^) that is closest to the center of the pericentromeric regions. When an assembly lacked a breakpoint because the centromere was fully assembled, we introduced an artificial breakpoint at the midpoint of the reference pericentromeric region. This adjustment ensures that all the assemblies for a given chromosome are aligned and anchored by their two ends, with only the longest assembly remaining unmodified.

### TE annotation of the 91 genomes

To annotate TEs that are present in each of the 91 genomes, we used RepeatMasker ^59^ and either the TAIR12 or the pan TE library with the options: -s -a -gff -no_is -nolow -cutoff 300. As RepeatMasker tends to annotate interstitial telomeric repeats, telomeric repeats, rDNA sequences, and other non-TE repetitive sequences as TEs, we used TRASH ^60^ and TRF ^61^ to identify them and removed them from the annotation. Specifically, sequences identified using TRASH and TRF were merged when closer than 200bp from each other and then masked when the merged sequence was larger than 1 kb.

### Detection and analysis of TE insertions polymorphisms (TIPs)

TIPs were identified using GraffiTE, a Nextflow-based workflow for the detection and analysis of TIPs in genome assemblies ^27^. GraffiTE was run from the main branch at Git revision f37674bd5fcd9e92e08a470911813a1c5807d084, using the 91 genome assemblies and either the pan TE or the TAIR12 TE libraries. The workflow was executed with Nextflow ^62^ version 25.10.2. TIPs detected using the pan TE or TAIR12 TE libraries were deemed identical if a) they were detected by both libraries within less than 10bp of each other, b) they belong to the same superfamily and c) they are detected in a mostly overlapping (90%) set of genomes.

### Generation of *ddm1-67* mutants

Two single-guide RNAs with similar efficiencies (sg6 and sg7, Fig. 4c) were designed using crispor ^63^ (https://crispor.gi.ucsc.edu/) to target the exons 6 and 7 of *DDM1* within a conserved region (based on 1001Genomes SNPs) of the protein ^64^. The DNA sequence for the two single-guide RNAs with their U6 promoters was synthesized by ThermoFisher (GeneArt) within the pDONR221 plasmid and then introduced by LR Gateway recombination into a binary-vector pDe-CAS9-DsRed plasmid carrying a Spectinomycin-resistance cassette. Integrity of the plasmid was verified by sequencing prior to it being introduced into *Agrobacterium tumefaciens* strain GV3101, which was then used to transform WT plants of each strain by floral dip following standard protocols.

Transformant T1 seeds were screened for DsRed fluorescence before PCR-genotyping seedling using primers amplifying 634bp from the U6 promoter of sg6 to the region upstream of DsRed (CGGTGCTTTTTTTGGATCC and TGCTTCTTGCTGAGCTCACA). Non-fluorescent T2 seeds from transgenic T1s were selected for PCR-verification of the absence of the transgene (no amplification with the primers indicated above) and for the presence of the *DDM1* deletion, using for the latter primers in exon 6 and intron 7 (CACGCCTTCCATCAATGCAA and AGCTGCCACCAGTGTTAACA, respectively). Non-transgenic individuals putatively hemizygous for the *DDM1* deletion were selected and verified by Sanger sequencing (Eurofins) of the WT- and deletion-size amplification products. These individuals were selfed to identify first-generation homozygous *ddm1-67* (*d/d*) and WT (*D/D*) offspring and these were selfed for one generation to obtain G2 individuals.

### Long read RNA-seq of *ddm1-67* mutant lines

RNA was extracted from from two young leaves collected from 15 second-generation *ddm1-67* individuals, 24 days after sowing, using Trizol (Invitrogen) and long-read RNA-seq was performed by GenomiqueENS (https://genomique.biologie.ens.fr/). Specifically, cDNAs were prepared using the SMART-Seq mRNA kit (Takara). Libraries were prepared using the cDNA-PCR Sequencing V14-Barcoding SQK-PCB114 and PCR Expansion EXP-PCA001 kits (Oxford Nanopore Technologies), before loading onto a PromethION R10.4.1 Flowcell (Oxford Nanopore Technologies). Read quality was assessed using FastQC (www.bioinformatics.babraham.ac.uk/projects/fastqc/) which was also used to inspect adapter-related sequence content. Adapter sequences were subsequently removed using cutadapt ^65^. Then the reads were mapped to TAIR12 genome assembly using minimap2 -ax splice ^47^.

### Whole Genome Bisulfite Sequencing of *ddm1-67* mutant lines

DNA was extracted using CTAB from the whole rosette of 12 of the 15 *ddm1-67* individuals used for RNA extraction as well as of 12 WT counterparts that were grown at the same time. Whole-genome bisulfite sequencing (WGBS-seq) was performed and analyzed as previously described ^15^. Methylation levels of TEs annotated with RepeatMasker and the pan TE library in each of the corresponding WT genome assembly were calculated separately for CG, CHG and CHH and averaged over each TE.

### Sequencing and de novo assembly of the genome of the five *ddm1-67* mutant lines

Sequencing and de novo assembly of the second-generation *ddm1-67* mutant line obtained for the Col-0, Cvi-0, C24, Ct-1 and Bd-0 strains was carried out on as for the 91 genomes (see above), except that for the use of the ONT MinION flow cell R10.4.1 instead of R9.4.1 and base calling with dorado (https://github.com/nanoporetech/dorado version 0.9.1; with dna_r10.4.1_e8.2_400bps_hac@v5.2.0) for the last four mutant strains. The absence of the transgene sequence (Supplementary File S1) was confirmed by BLAST.

Putative off-target sites (up to 5 mismatches allowed) for both single guide RNAs were computed using the Cas-OFFinder tool ^36^, where the reference genome used is that of the corresponding WT genome assembly. The sequence of each putative off-target site was manually compared between the reference and mutant genome and no off-targets were found.

### Detection of transposition events within long-reads

Transposition events were detected using the OUTSIDER module of the tool TrEMOLO ^35^, version 2.4, installed through Singularity. Parameters and settings were those provided in the TrEMOLO configuration file available on GitHub, with the exception that SVIM ^66^ was used for the detection of outsider structural variants. The resulting TE_INFOS.bed file was subsequently filtered to only retain high-confidence transposition events, i.e. those with a target site duplication (TSD, except for *Helitron,* which do not produce TSDs upon insertion), and that are located in regions with a sequencing coverage below 100×. In addition, the size of the outsider was required to be consistent with that of the corresponding structural variant detected by SVIM (70%–130% of the outsider size), to avoid possible composite events (e.g. pack-MULE-like structures) or local recombination events encompassing a TE. Finally, outsiders were required to cover at least 80% of the length of the corresponding consensus sequence (psize_TE). The results obtained for the 91 WT genomes were further filtered to only retain outsiders covered by just one or two reads, to avoid false positives due to residual heterozygosity. As DNA was extracted from pools of 5-10 sibling individuals, a TE insertion with such low coverage likely reflects somatic transposition, or else germline transposition inherited by just one individual of the pool, rather than residual heterozygosity. Additionally, because sequence coverage for the 91 WT genomes varied greatly (10 to 68X), we normalized the number of outsiders detected in each of them by 10X coverage.

### PCR validation of outsiders

The 17 outsiders covered by two reads in the 13 individuals for which we had genomic DNA used for genome sequencing, were chosen for PCR validation. Three primers were designed for each outsider, two that flank the insertion, and one at an extremity of the putative insertion. PCRs were run with both flanking primers to amplify the empty site, or with the insertion primer and each of the flanking primers to amplify the junction, either left or right, between the insertion and the flanking sequence. We could not obtain any amplification of the empty side for one outsider tested in this manner. Among the other 16 outsiders, 13 were confirmed by PCR.

### Identification of donor copies

The sequence of outsiders identified by TrEMOLO in the five *ddm1-67* mutant lines was extracted from the corresponding reads and compiled into multi-FASTA files with the sequence of the longest copies of the corresponding TE family that are present in the WT genome assembly. After MAFFT ^52^ alignment (with --reorder and --adjustdirectionaccurately) of these sequences, we built the NJ trees with Biopython ^67^ and visualized them using ETE 3 ^68^. Putative donor copies were identified by combining information obtained from the NJ trees, using as a proxy of genetic distance the number of SNPs and indels between each outsider and each annotated copy. In some cases, the NJ tree topologies did not provide sufficient resolution, or the genetic distances between candidate copies were too similar to unambiguously assign a donor copy.

### Estimation of the population frequency of donor copies

From the coordinates of the donor copy in its original genome, we selected the three closest genes upstream and downstream with bedtools closest ^69^, and asked in all other 90 genome assemblies if a TE copy from the same TE family is present at the same location, together, with at least two of the three genes on either side. Visual inspection was used to verify that the TE copy is present at the expected position in every case. Were this not the case, we excluded the donor copy from our population frequency analysis.

### Estimation of the age of donor copies

For the *ATMU3* and *ATCOPIA74* donor copies identified respectively on chromosome 1 and 4 of C24, we created multiple sequence alignments across the carrier genomes using MAFFT ^52^(--auto), and calculated pairwise nucleotide differences using Biopython ^67^. Age was then estimated by taking the highest of 84 SNPs between any two copies and using the Single-Nucleotide Mutation (SNM) rate calculated for TEs (1.4E-8 per site per generation, ^70^).

### Phylogenetic and pairwise diversity analysis of the regions surrounding the high population frequency *ATMU3* and *ATCOPIA74* insertions

For the two insertions, we extracted from all 91 genomes the sequence that contains the closest upstream and downstream genes. The insertion itself was removed from that sequence in all strains where it is present in order to generate a multiple alignment (MAFFT ^52^ --auto), which was used to build a ML tree with IQ-TREE 2 ^56^, visualized using iTol ^55^.

Nucleotide diversity of carriers and non-carriers of the *ATCOPIA74* chromosome 4 insertion was calculated using bi-allelic SNPs from the 1,001 Genomes with a minimum missing genotyping rate of 0.2 and located on chromosome 4. Pairwise diversity was calculated in consecutive 5kb windows using selscan -v 1.2.0a ^71^ following phasing and inference using shapeit v2.r904 ^72^.

Moving averages were calculated over 20 consecutive 5kb windows for whole chromosome visualization, or over 2 consecutive windows for visualization of the first 300 kb region containing the *ATCOPIA74* insertion.

### Miscellaneous

All plots and panels of the figures were generated using matplotlib ^73^ or ggplot2 (ggplot2.tidyverse.org/). Tukey’s honestly significant difference (HSD) test, as implemented in the statsmodels Python package with pairwise_tukeyhsd (https://www.statsmodels.org/), was used to assess the statistical significance of the difference of the number of bases annotated as TEs and of the number of full-length TE copies identified between genetic groups (Fig. 2 and 3). Wilcoxon tests were performed in R to assess the difference of methylation levels over TEs between WT and *ddm1-67* mutant lines (Fig. 4).

## Supplementary Datasets

Supplementary File S1 - Phylogenies of consensus of all the superfamilies, coming from pantera, TAIR12 TE library, RepetDB TE library or Repbase database.

Supplementary File S2 - pan TE library fasta sequences

Supplementary data S1 - TE insertion polymorphisms detected by GraffiTE with TAIR12 TE library

Supplementary data S2 - TE insertion polymorphisms detected by GraffiTE with our pan TE library

Supplementary data S3 - All 509 filtered outsiders detected by TrEMOLO across the 91 genome assemblies

Supplementary data S4 - All 108 filtered outsiders detected by TrEMOLO across the five *ddm1-67* mutant genome assemblies.

Supplementary data S5 - All donor copies identified across the five *ddm1-67* mutant genome assemblies (with information about allele frequency, etc…)

Supplementary Table S1: Assembly stats (BUSCO completeness, N50 reads, N50 contigs, assembly length), project name in NCBI,…

Supplementary Table S2: recap of all the pantera TEs, trimming (TEtrimmer automatic or GUI), superfamily attributed by RepeatClassifier, closest reference TE, match stats (identity, query cov, target cov), new names

Supplementary Table S3: TE families with experimental evidence of mobility (with refs), and their corresponding pantera sequence if found.

Supplementary Table S4: list of 489 TE families from TAIR12, and 5 RepBase, not found by pantera

Supplementary Table S5: list of Primers for PCR, coordinates of the insertions.

## Data Availability

ONT-seq read-data (fastq), *de novo* genome assemblies (fasta) of 71 natural strains, pangenome graph alignment (gfa) and TE annotation (gff) of 91 genomes, as well as raw BS-seq, ONT RNA-seq, ONT-seq read-data (fastq), and *de novo* genome assemblies (fasta) of *ddm1-67* mutant lines generated and analyzed for this study are available in the European Nucleotide Archive (ENA) under project PRJEBXXXX.

WGS data was obtained in FASTQ format for the 1047 strains of the 1001genomes.org project from NCBI SRA Project PRJNA273563. Raw RNA-seq data from the 1001 Genomes was obtained from NCBI GEO Accession GSE80744.

## Acknowledgments

We thank Richard Durbin for hosting AP in his group and introducing him to pantera. We also thank Richard Durbin and Pío Sierra for sharing early versions of pantera and for discussions. We thank the 1001Genomes Plus consortium, especially Detlef Weigel and Magnus Nordborg, as well as Leandro Quadrana (IPS2, Orsay), and members of the Baduel group for their inputs. Computational analyses were performed on the IBENS bioclust computing cluster and the Institut Français de Bioinformatique (IFB) cluster. We thank the IT & bioinformatics platforms, and especially Alexandra Louis for their support. We thank Fabien Nogué (IJPB, INRAE Versailles) for sharing DeCas9 plasmid and for his advice on the design of guide RNAs, as well as Abdou Khadre Diouf for preliminary analyses of the genome assemblies.

## Authors contributions

VC and PB designed the study, supervised the work, and wrote the manuscript together with AP. AP performed most of the bioinformatic analyses, with additional contributions from MR, NAG and LDO. EC performed DNA and ONT-sequencing extractions for the 71 strains (including the 5 *ddm1* mutant strains) and RNA extractions for *ddm1-67* mutants, with additional support from MR. The 5 *ddm1-67* mutant strains were generated by PB, CX, CC and DS.

All authors read and approved the manuscript.

## References

1. Chuong, E. B., Elde, N. C. & Feschotte, C. Regulatory activities of transposable elements: from conflicts to benefits. Nat Rev Genet 18, 71–86 (2017).

2. Rech, G. E. et al. Population-scale long-read sequencing uncovers transposable elements associated with gene expression variation and adaptive signatures in Drosophila. Nat Commun 13, 1948 (2022).

3. Tsay, Y. F., Frank, M. J., Page, T., Dean, C. & Crawford, N. M. Identification of a mobile endogenous transposon in Arabidopsis thaliana. Science 260, 342–344 (1993).

4. Singer, T., Yordan, C. & Martienssen, R. A. Robertson’s Mutator transposons in A. thaliana are regulated by the chromatin-remodeling gene Decrease in DNA Methylation (DDM1). Genes Dev 15, 591–602 (2001).

5. Miura, A. et al. Mobilization of transposons by a mutation abolishing full DNA methylation in Arabidopsis. Nature 411, 212–214 (2001).

6. Kato, M., Miura, A., Bender, J., Jacobsen, S. E. & Kakutani, T. Role of CG and non-CG methylation in immobilization of transposons in Arabidopsis. Curr Biol 13, 421– 426 (2003).

7. Lippman, Z. et al. Role of transposable elements in heterochromatin and epigenetic control. Nature 430, 471–476 (2004).

8. Tsukahara, S. et al. Bursts of retrotransposition reproduced in Arabidopsis. Nature 461, 423–426 (2009).

9. Mirouze, M. et al. Selective epigenetic control of retrotransposition in Arabidopsis. Nature 461, 427–430 (2009).

10. Fu, Y. et al. Mobilization of a plant transposon by expression of the transposon-encoded anti-silencing factor. EMBO J 32, 2407–2417 (2013).

11. Quadrana, L. et al. Transposition favors the generation of large effect mutations that may facilitate rapid adaption. Nat. Commun. 10, 3421 (2019).

12. Quadrana, L. et al. The Arabidopsis thaliana mobilome and its impact at the species level. Elife 5, (2016).

13. Baduel, P. et al. Genetic and environmental modulation of transposition shapes the evolutionary potential of Arabidopsis thaliana. Genome Biol 22, 138 (2021).

14. Sierra, P. & Durbin, R. Identification of transposable element families from pangenome polymorphisms. Mob DNA 15, 13 (2024).

15. Baduel, P. et al. Transposable elements are vectors of recurrent transgenerational epigenetic inheritance. Science 390, eady3475 (2025).

16. Alonso-Blanco, C. et al. 1,135 Genomes Reveal the Global Pattern of Polymorphism in Arabidopsis thaliana. Cell 166, 481–491 (2016).

17. Bao, W., Kojima, K. K. & Kohany, O. Repbase Update, a database of repetitive elements in eukaryotic genomes. Mob DNA 6, 11 (2015).

18. Tegenfeldt, F. et al. OrthoDB and BUSCO update: annotation of orthologs with wider sampling of genomes. Nucleic Acids Res 53, D516–D522 (2025).

19. Lian, Q. et al. A pan-genome of 69 Arabidopsis thaliana accessions reveals a conserved genome structure throughout the global species range. Nat Genet 56, 982–991 (2024).

20. Igolkina, A. A. et al. A comparison of 27 Arabidopsis thaliana genomes and the path toward an unbiased characterization of genetic polymorphism. Nat. Genet. 57, 2289– 2301 (2025).

21. Garrison, E. et al. Building pangenome graphs. Nature Methods 21, 2008–2012 (2024).

22. Amselem, J. et al. RepetDB: a unified resource for transposable element references. Mob DNA 10, 6 (2019).

23. Flynn, J. M. et al. RepeatModeler2 for automated genomic discovery of transposable element families. Proc Natl Acad Sci U S A 117, 9451–9457 (2020).

24. Alonso-Blanco, C. C., et al. The 1001G+ project: A curated collection of *Arabidopsis thaliana* long-read genome assemblies to advance plant research. bioRxiv (2024) doi:10.1101/2024.12.23.629943.

25. Koch, M. A., Haubold, B. & Mitchell-Olds, T. Comparative evolutionary analysis of chalcone synthase and alcohol dehydrogenase loci in Arabidopsis, Arabis, and related genera (Brassicaceae). Mol Biol Evol 17, 1483–1498 (2000).

26. Stuart, T. et al. Population scale mapping of transposable element diversity reveals links to gene regulation and epigenomic variation. Elife 5, e20777 (2016).

27. Groza, C., Chen, X., Wheeler, T. J., Bourque, G. & Goubert, C. A unified framework to analyze transposable element insertion polymorphisms using graph genomes. Nat Commun 15, 8915 (2024).

28. Raingeval, M. et al. Retrotransposon-driven environmental regulation of FLC leads to adaptive response to herbicide. Nat. Plants 10, 1672–1681 (2024).

29. Kawakatsu, T. et al. Epigenomic Diversity in a Global Collection of Arabidopsis thaliana Accessions. Cell 166, 492–505 (2016).

30. Wu, D. et al. The prodomain of Arabidopsis metacaspase 2 positively regulates immune signaling mediated by pattern-recognition receptors. New Phytol 241, 430–443 (2024).

31. Coll, N. S. et al. Arabidopsis type I metacaspases control cell death. Science 330, 1393–1397 (2010).

32. Aoun, N. et al. Natural variation in an NLR pair confers thermostable resistance to a devastating bacterial pathogen. bioRxiv (2025) doi:10.64898/2025.12.15.694456.

33. Todesco, M. et al. Natural allelic variation underlying a major fitness trade-off in Arabidopsis thaliana. Nature 465, 632–636 (2010).

34. Lyons, R. et al. Investigating the Association between Flowering Time and Defense in the Arabidopsis thaliana-Fusarium oxysporum Interaction. PLoS One 10, e0127699 (2015).

35. Mohamed, M. et al. TrEMOLO: accurate transposable element allele frequency estimation using long-read sequencing data combining assembly and mapping-based approaches. Genome Biol 24, 63 (2023).

36. Bae, S., Park, J. & Kim, J.-S. Cas-OFFinder: a fast and versatile algorithm that searches for potential off-target sites of Cas9 RNA-guided endonucleases. Bioinformatics 30, 1473–1475 (2014).

37. Zemach, A. et al. The Arabidopsis nucleosome remodeler DDM1 allows DNA methyltransferases to access H1-containing heterochromatin. Cell 153, 193–205 (2013).

38. Movilli, A., Sushko, S., Rabanal, F. A. & Weigel, D. Long-read detection of transposable element mobilization in the soma of hypomethylated Arabidopsis thaliana individuals. Genome Biol 26, 231 (2025).

39. Fulgione, A. et al. Parallel reduction in flowering time from de novo mutations enable evolutionary rescue in colonizing lineages. Nat Commun 13, 1461 (2022).

40. Scarpa, A. et al. Double trouble: two retrotransposons triggered a cascade of invasions in Drosophila species within the last 50 years. Nat Commun 16, 516 (2025).

41. Pianezza, R., Scarpa, A., Haider, A., Signor, S. & Kofler, R. Spatiotemporal Tracking of Three Novel Transposable Element Invasions in Drosophila melanogaster over the Last 30 Years. Mol Biol Evol 42, (2025).

42. Platt, A. et al. The scale of population structure in Arabidopsis thaliana. PLoS Genet 6, e1000843 (2010).

43. Alonge, M. et al. Automated assembly scaffolding using RagTag elevates a new tomato system for high-throughput genome editing. Genome Biol 23, 258 (2022).

44. Weigel, D. & Mott, R. The 1001 genomes project for Arabidopsis thaliana. Genome Biol 10, 107 (2009).

45. Naish, M. et al. The genetic and epigenetic landscape of the centromeres. Science 374, eabi7489 (2021).

46. Ahsan, M. U., Liu, Q., Fang, L. & Wang, K. NanoCaller for accurate detection of SNPs and indels in difficult-to-map regions from long-read sequencing by haplotype-aware deep neural networks. Genome Biol 22, 261 (2021).

47. Li, H. Minimap2: pairwise alignment for nucleotide sequences. Bioinformatics 34, 3094–3100 (2018).

48. Pisupati, R. et al. Verification of Arabidopsis stock collections using SNPmatch, a tool for genotyping high-plexed samples. Sci Data 4, 170184 (2017).

49. Chang, C. C. et al. Second-generation PLINK: rising to the challenge of larger and richer datasets. Gigascience 4, 7 (2015).

50. Lamesch, P. et al. The Arabidopsis Information Resource (TAIR): improved gene annotation and new tools. Nucleic Acids Res 40, D1202–10 (2012).

51. Shumate, A. & Salzberg, S. L. Liftoff: accurate mapping of gene annotations. Bioinformatics 37, 1639–1643 (2021).

52. Katoh, K. & Standley, D. M. MAFFT multiple sequence alignment software version 7: improvements in performance and usability. Mol Biol Evol 30, 772–780 (2013).

53. Capella-Gutiérrez, S., Silla-Martínez, J. M. & Gabaldón, T. trimAl: a tool for automated alignment trimming in large-scale phylogenetic analyses. Bioinformatics 25, 1972–1973 (2009).

54. Borowiec, M. L. AMAS: a fast tool for alignment manipulation and computing of summary statistics. PeerJ 4, e1660 (2016).

55. Letunic, I. & Bork, P. Interactive Tree of Life (iTOL) v6: recent updates to the phylogenetic tree display and annotation tool. Nucleic Acids Res 52, W78–W82 (2024).

56. Minh, B. Q. et al. IQ-TREE 2: New Models and Efficient Methods for Phylogenetic Inference in the Genomic Era. Mol Biol Evol 37, 1530–1534 (2020).

57. Qian, J. et al. TEtrimmer: a tool to automate the manual curation of transposable elements. Nat Commun 16, 8429 (2025).

58. Wlodzimierz, P. et al. Cycles of satellite and transposon evolution in Arabidopsis centromeres. Nature 618, 557–565 (2023).

59. Arian FA Smit, Robert Hubley, P Green. RepeatMasker. RepeatMasker Open-4.0. 2013–2015 http://www.repeatmasker.org.

60. Wlodzimierz, P., Hong, M. & Henderson, I. R. TRASH: Tandem Repeat Annotation and Structural Hierarchy. Bioinformatics 39, (2023).

61. Benson, G. Tandem repeats finder: a program to analyze DNA sequences. Nucleic Acids Res 27, 573–580 (1999).

62. Di Tommaso, P. et al. Nextflow enables reproducible computational workflows. Nat Biotechnol 35, 316–319 (2017).

63. Concordet, J.-P. & Haeussler, M. CRISPOR: intuitive guide selection for CRISPR/Cas9 genome editing experiments and screens. Nucleic Acids Res 46, W242– W245 (2018).

64. Jeddeloh, J. A., Stokes, T. L. & Richards, E. J. Maintenance of genomic methylation requires a SWI2/SNF2-like protein. Nat. Genet. 22, 94–97 (1999).

65. Martin, M. Cutadapt removes adapter sequences from high-throughput sequencing reads. EMBnet J. 17, 10 (2011).

66. Heller, D. & Vingron, M. SVIM: structural variant identification using mapped long reads. Bioinformatics 35, 2907–2915 (2019).

67. Cock, P. J. A. et al. Biopython: freely available Python tools for computational molecular biology and bioinformatics. Bioinformatics 25, 1422–1423 (2009).

68. Huerta-Cepas, J., Serra, F. & Bork, P. ETE 3: Reconstruction, Analysis, and Visualization of Phylogenomic Data. Mol Biol Evol 33, 1635–1638 (2016).

69. Quinlan, A. R. & Hall, I. M. BEDTools: a flexible suite of utilities for comparing genomic features. Bioinformatics 26, 841–842 (2010).

70. Weng, M.-L. et al. Fine-Grained Analysis of Spontaneous Mutation Spectrum and Frequency in. Genetics 211, 703–714 (2019).

71. Torres, R., Szpiech, Z. A. & Hernandez, R. D. Human demographic history has amplified the effects of background selection across the genome. PLoS Genet 14, e1007387 (2018).

72. O’Connell, J. et al. A general approach for haplotype phasing across the full spectrum of relatedness. PLoS Genet 10, e1004234 (2014).

73. The Matplotlib Development Team. Matplotlib: Visualization with Python. (Zenodo, 2024). doi:10.5281/ZENODO.14249941.

