## Supplementary Figures for "Fragmented landscapes of mobile transposable element families bolster the diversity of large-effect alleles in Arabidopsis"

Running title: Extensive diversity of mobile transposable elements in Arabidopsis

Aurélien Petit[1], Erwann Caillieux[1], Mathilde Reignier[1], Ndeye Amy Gueye[1], Louna De Oliveira[1], Ciana Xu[1], Clio Cordier[1], Dersu Sezginmert[1], Vincent Colot[1*], Pierre Baduel[1*]

[1] *Institut de Biologie de l'École Normale Supérieure, Université Paris Sciences et Lettres (PSL), UMR 8197 CNRS, U1024 INSERM, 46 rue d'Ulm, 75005 Paris, France*

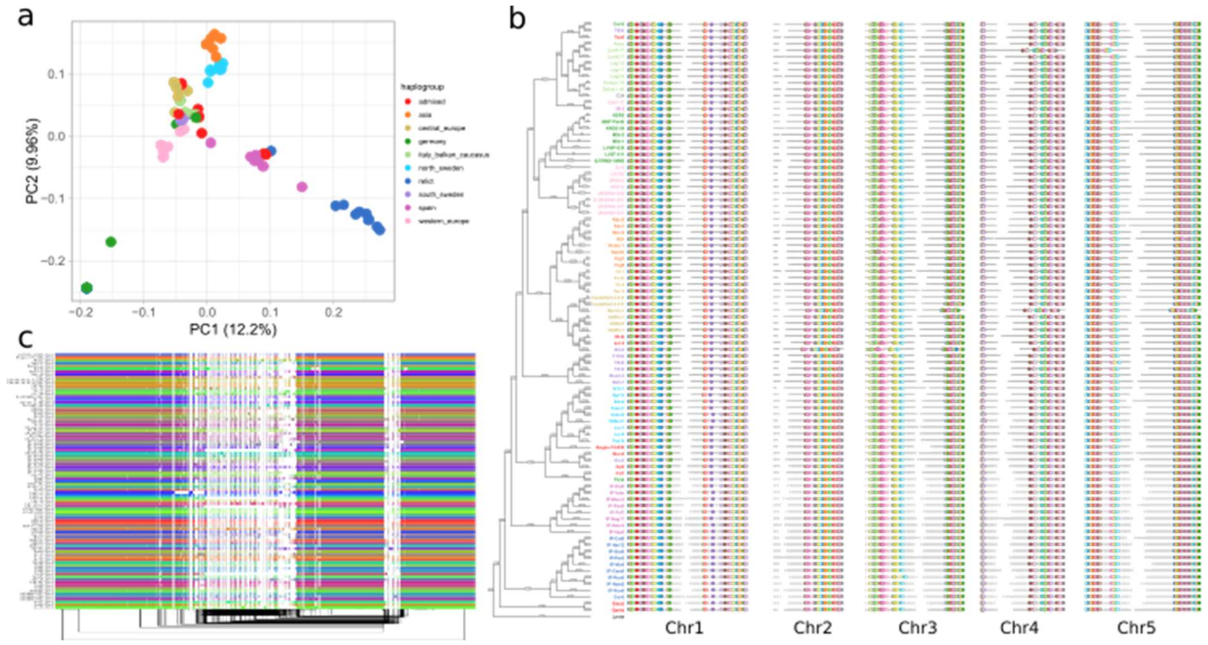


**Figure S1. Relatedness of the 91 genome assemblies**

a) PCA of the 91 genomes using SNPs obtained from the 1001 genomes project. b) Alignment by chromosome ends (see Methods). Randomly selected genes from annotations transferred with LiftOff (see Methods) are highlighted in colored circles to highlight the conservation of synteny within chromosome arms. c) Pan-genome graph of chromosome 1 produced by pggb. Structural variants are not colored in the all-to-all alignment and are highlighted with black lines.

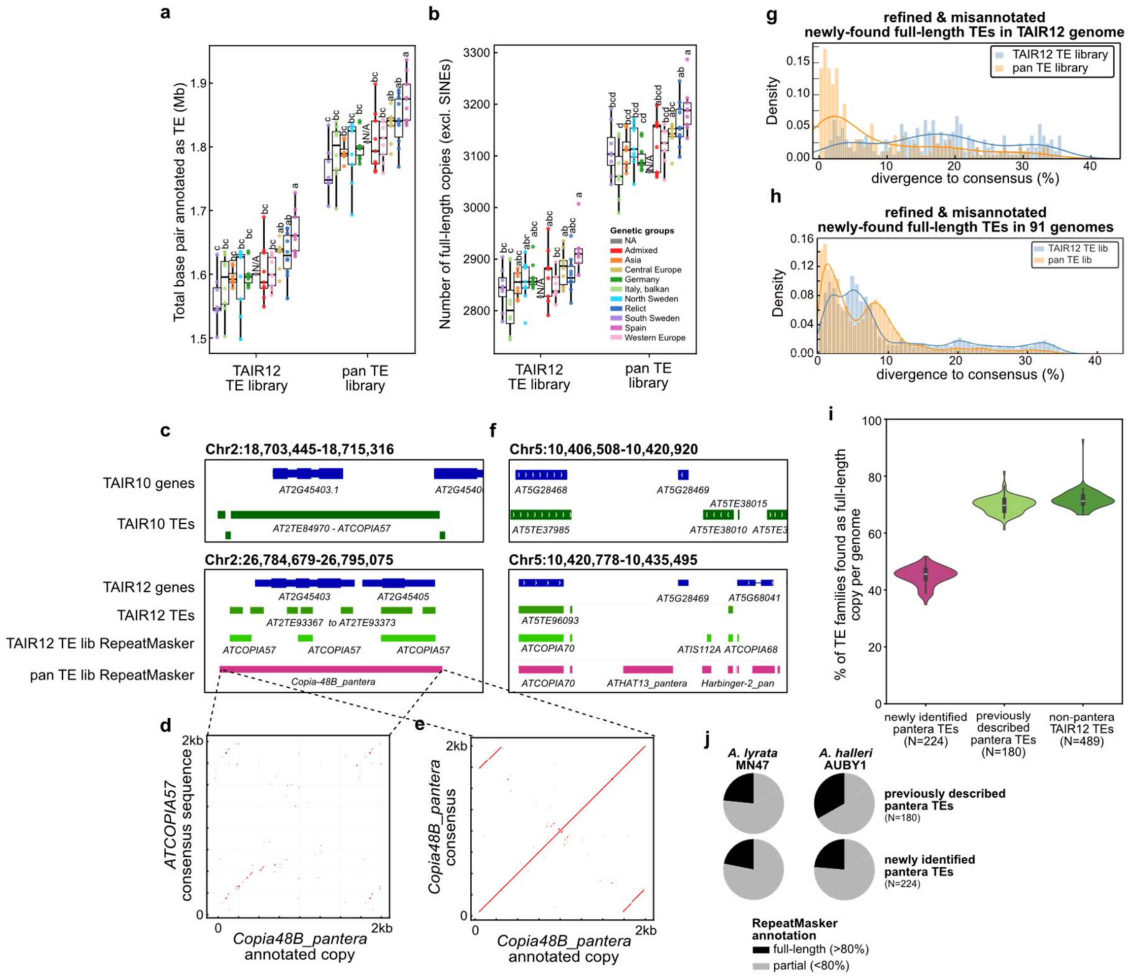


**Figure S2. Improvement of TE annotation using the pan TE library**

a) Total TE content and b) total number of full-length copies (excluding SINEs) per genome based on TE annotation using RepeatMasker and either the TAIR12 or the pan TE library. Strains are collapsed by genetic groups and letters indicate post-hoc Tukey test groups, as in Fig. 2. c) Comparison of TE annotations present over the ~2kb region of Chr2 in TAIR10 (top) and TAIR12 (bottom) surrounding the newly found *Copia-48B* full-length copy annotated using the pan TE library. The newly found *Copia-48B* was previously annotated as one (TAIR10) or multiple partial copies of *ATCOPIA57* (TAIR12). d) and e) Dotplots of the *Copia-48B* annotated copy against the *ATCOPIA57* and *Copia-48B_pantera* consensus, respectively. f) Comparison of TE annotations present over the ~14kb region of Chr5 in TAIR10 (top) and TAIR12 (bottom) surrounding the newly found *ATHAT13* full-length copy annotated using the pan TE library. The *ATHAT13* copy was previously unannotated in both TAIR10 and TAIR12, as well as using the TAIR12 TE library. g) Divergence to TAIR12 or pan TE consensus (as given by RepeatMasker) of copies annotated in the TAIR12 genome assembly as full-length using the pan TE library but as partial using the TAIR12 TE library. h) Divergence to TAIR12 or pan TE consensus of copies annotated in 91genome assemblies as full-length using the pan TE library but as partial using the TAIR12 TE library.

The presence of two peaks of divergence, around 0-1% and ~9% suggests the existence of TE family sublineages. i) Percentage of TE families from each of the three categories (newly identified by pantera, previously-described and not identified by pantera), that have at least one full-length copy per genome. j) Proportion of previously or newly identified TE families that are annotated as full-length (>80%) or partial (<80%) in the *MN47 A. lyrata* or *AUBY1 A. halleri* genomes.

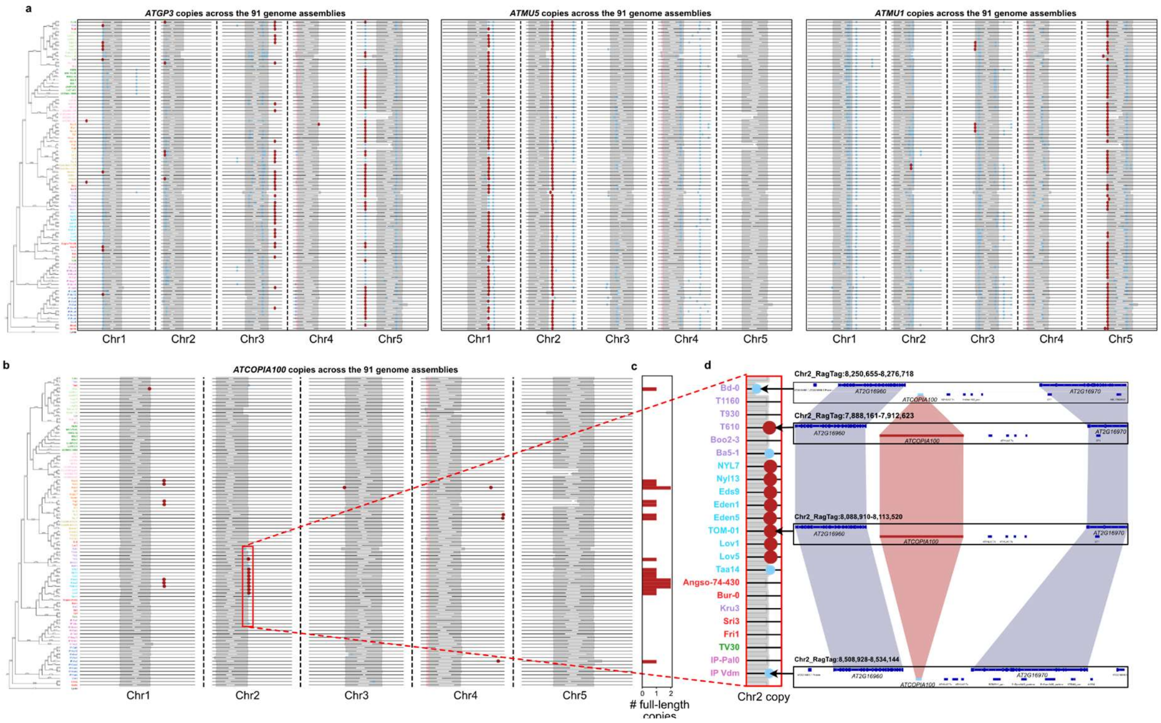


**Figure S3 Example of a TE copy present at the same location either as a full-length or partial copy across genetic groups.**

a) Full-length (red) and partial (blue) copies of *ATGP3* (left), *ATMU5* (middle), and *ATMU1* (right) present in the 91 genomes, aligned from their chromosome ends and ordered according to their phylogenetic relationship. Genetic groups are colored as in Fig. 1. b) Full-length (red) and partial (blue) copies of *ATCOPIA100* present in the 91 genomes, aligned from their chromosome ends and ordered according to their phylogenetic relationship. Genetic groups are colored as in Fig. 1. c) Number of *ATCOPIA100* full-length copies per genome. d) Genome browser view of four genomes carrying the *ATCOPIA100* copy located on chromosome 2, either as full-length or else as a solo-LTR. Note that the IP-Vdm genome, which carries a solo-LTR and all other genomes from the Spain genetic groups appear to be devoid of any full-length *ATCOPIA100* copy.

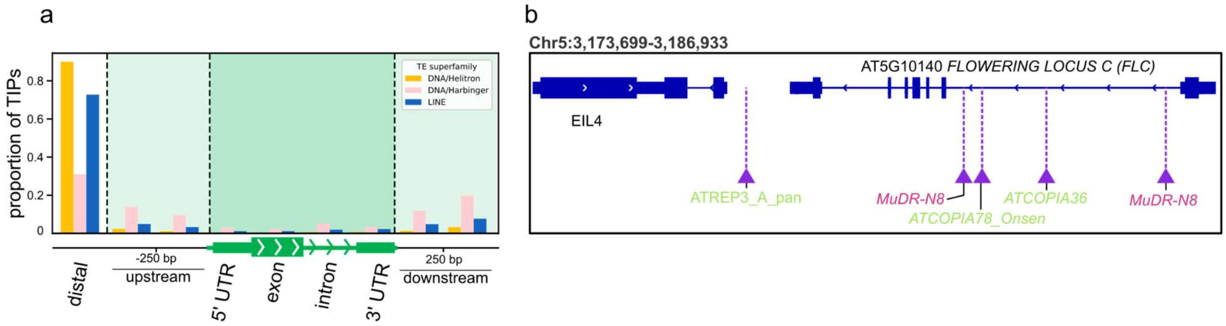


**Figure S4. Position of TIPs relative to genes**

a) Proportion of *Helitron*, *Harbinger,* and *LINE* TIPs detected using the pan TE library according to their position relative to genes. b) Genome browser view of the *FLC* locus indicating the position and identity of the five TIPs detected over it across the 91 genomes when using the pan TE library. TIPs not previously detected are highlighted in pink and in green otherwise.

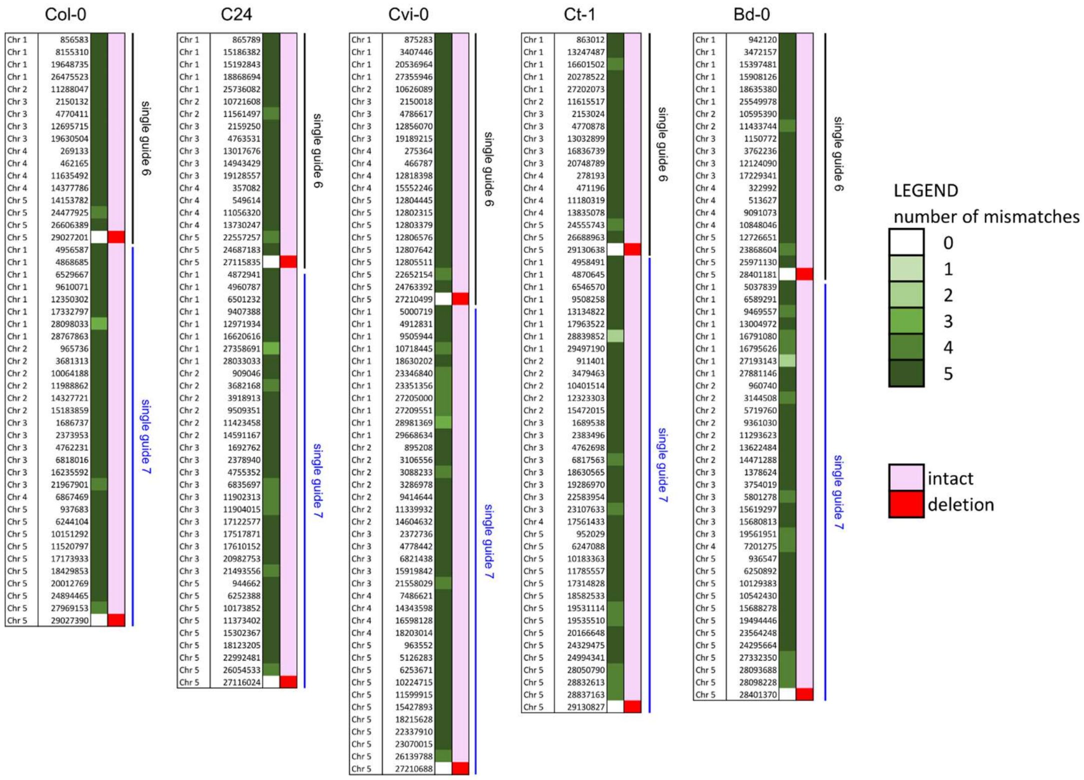


**Figure S5: Absence of predicted off-targeting for the generation of the *ddm1-67* deletion**

Position of predicted targets with up to five mismatches for each guide RNA and each genome. Number of mismatches are indicated in shades of green. The only perfect matches are in exons 6 and 7 of *DDM1* and are in white. Visual inspection of all predicted targets and surrounding sequences revealed no mutation (pink), except for the expected deletion over the *DDM1* locus (red).

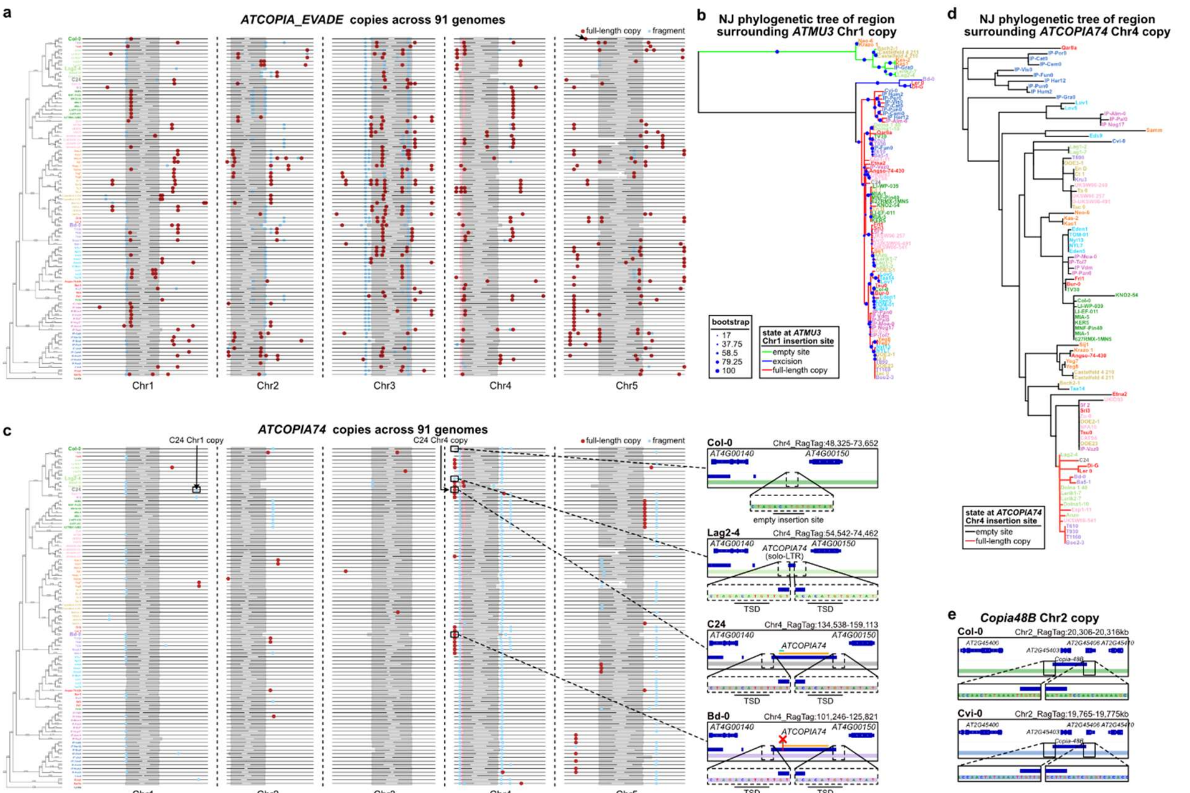


**Figure S6: Evolutionary trajectories of the mobile *ATCOPIA74* and *Copia-48B* insertions**
a) Position of the *ATCOPIA_EVADE* partial (blue) and full-length (red) copies present in the 91 genomes, aligned from their chromosome ends and ordered according to their phylogenetic relationship. Genetic groups are colored as in Fig. 1. b) Neighbor-joining phylogenetic tree of the region surrounding the chromosome 1 donor *ATMU3* copy detected in the C24 *ddm1-67* mutant line, for the 91 genomes, after removing the sequence of the copy in the carriers. Each branch is highlighted based on the allele of a given strain (i.e. full-length copy in red, excision footprint in blue and absence allele in green). c) Position of the *ATCOPIA74* partial (blue) and full-length (red) copies present in the 91 genomes, aligned from their chromosome ends and ordered according to their phylogenetic relationship. Genetic groups are colored as in Fig. 1. The region carrying the donor copy of Chr4 identified in the C24 *ddm1-67* mutant line is highlighted on the right panels for four strains, which either carry the donor copy as full-length (C24 and Bd-0, with a red cross to indicate the non-sense substitution) or a partial copy (Lag2-4) or an empty site (Col-0). Sequences at the insertion site are shown. Empty site and TSDs are underlined. Open reading frame (ORF) and endonuclease domain are annotated in orange and cyan respectively, with the nonsense substitution (red cross) present in the Bd-0 insertion. d) Neighbor-joining phylogenetic tree of the region surrounding the chromosome 4 donor *ATCOPIA74* copy detected in the C24 *ddm1-67* mutant line, for the 91 genomes, after removing the sequence of the copy in the carriers. Each branch is highlighted based on the allele of a given strain (i.e. full-length copy in red and absence allele in black). e) Genome browser view of the region around the chromosome 2 donor *Copia-48B* copy detected in the Col-0 *ddm1-67* mutant line, in the Col-0 (top panel) or in the Cvi-0 genome assembly (bottom panel). Sequences at the left and right junctions are shown.
